# 3D confinement regulates stem cell fate

**DOI:** 10.1101/2021.05.02.442094

**Authors:** Oksana Y. Dudaryeva, Aurelia Bucciarelli, Giovanni Bovone, Shibashish Jaydev, Nicolas Broguiere, Marwa al-Bayati, Marco Lütolf, Mark W. Tibbitt

## Abstract

Biophysical properties of the cellular microenvironment, including stiffness and geometry, influence cell fate. Recent findings have implicated geometric confinement as an important regulator of cell fate determination. Our understanding of how mechanical signals direct cell fate is based primarily on two-dimensional (2D) studies. To investigate the role of confinement on stem cell fate in three-dimensional (3D) culture, we fabricated a single cell microwell culture platform and used it to investigate how niche volume and stiffness affect human mesenchymal stem cell (hMSC) fate. The viability and proliferation of hMSCs in confined 3D microniches were compared with the fate of unconfined cells in 2D culture. Physical confinement biased hMSC fate, and this influence was modulated by the niche volume and stiffness. The rate of cell death increased, and proliferation markedly decreased upon 3D confinement. We correlated the observed differences in hMSC fate to YES-associated protein (YAP) localization. In 3D microniches, hMSCs displayed primarily cytoplasmic YAP localization, indicating reduced mechanical activation upon confinement. These results demonstrate that 3D geometric confinement can be an important regulator of cell fate, and that confinement sensing is linked to canonical mechanotransduction pathways.

## Introduction

Cell fate is orchestrated both by the internal machinery of the cell and through reciprocal interactions with the surrounding extracellular matrix (ECM).^1, 2^ Extracellular regulators of cell function include biochemical cues, such as growth factors and cytokines, and biophysical cues, such as niche elasticity, geometry, and topography.^3–5^ The mechanisms by which biophysical signals influence cell function have been elucidated primarily using 2D culture, including standard tissue culture polystyrene and micropatterned substrates. Seminal findings in 2D revealed several mechanisms by which biophysical cues direct cell fate.^6–11^ For example, matrix elasticity induced both short-term and latent effects on stem cell function and differentiation.^5, 12–14^ Substrate topography controlled cell morphology, orientation, and proliferation.^15,16^ Geometric confinement in 2D directed cell fate; decreased cell spread area led to decreased proliferation and increased cell death.^17,18^ In total, these observations indicate a robust relationship between the physical properties of the microenvironment, cell morphology, and cell function.

While 2D culture has been particularly useful for studying biological processes, it fails to recapitulate critical characteristics of the native 3D cell niche and limits our ability to understand how biophysical properties, such as geometric confinement and stiffness, regulate cell fate. Hydrogels have been engineered specifically as ECM mimics to investigate the influence of matrix properties on cell function in 3D.^19–21^ Despite the broad utility of 3D platforms, standard encapsulation in bulk materials offers limited control over the geometry and volume of the individual cell niches. To address this limitation, microfabricated platforms have been developed that control previously inaccessible aspects of the 3D cellular environment, including niche geometry and volume.^22,23^ Microfabricated hydrogel platforms demonstrated that cell morphology and volume in 3D influenced cell shape, contractility, and transcription factor activity.^24,25^ Many of these micropatterned platforms employed geometrically-defined wells that constrained cells to attain morphologies that deviated from unconfined 2D cell shape. The wells were often similar to the average cell volume and smaller than the average cell spread length, confining the cells relative to standard 2D culture. Recent studies have demonstrated that 3D confinement can impact cell behavior constraining common cellular processes, such as spreading, migration, and proliferation, which require changes in cell volume, shape, or movement.^26^ Furthermore, cellular responses to confinement have been suggested to vary with matrix stiffness.^27,28^

A careful investigation of how 3D geometric confinement and stiffness of the cell niche couple to influence cell fate requires simultaneous control over the mechanical and geometric parameters of the cell microenvironment. Therefore, we developed a tunable and micropatterned culture platform to study how single cell fate was governed by niche properties in 3D. We encapsulated single hMSCs in individual niches with varied volume and stiffness and quantified how these properties affected cell viability and proliferation. As an unconfined control, hMSCs were seeded on 2D hydrogels with equivalent mechanical properties. In confined 3D niches, the viability of the hMSCs was reduced at low stiffness, most notably in low volume niches. The proliferation rate of hMSCs in confined 3D environments decreased significantly relative to 2D. To investigate if these effects were related to known mechanotransduction pathways, we assessed YAP localization in all conditions. Nuclear localization of YAP in cells in 3D niches was low correlated with cell proliferation. This suggested that the influence of 3D confinement on stem cell fate is coordinated through canonical mechanosensitive signaling pathways.

## Results

### Single cell microniche arrays to investigate physical regulation of cell fate in 3D

To investigate the role of geometric confinement and niche elasticity on single stem cell fate in 3D, we fabricated arrays of microniches with controlled geometry, volume, and stiffness (**Figure 1a,b**). We used poly(ethylene glycol) (PEG) hydrogels as the base material for the platform (**Figure 1c**). To form the gel, 10 kDa eight-arm PEG macromers functionalized with norbornene (PEG-NB) were reacted with a dithiol (dithiothreitol; DTT) via a photoinitiated thiol–ene reaction.^29^ To enable cell adhesion, the inert PEG hydrogel was functionalized with thiol-containing peptides. Specifically, we included the ECM-derived adhesion peptide CRGDS, which engages integrins (e.g., αvβ1, αvβ3 and αvβ5) and enables cell adhesion and spreading.^30^ To create fully confined 3D single cell environments, we sealed the niches with a second, non-patterned hydrogel layer using an enzymatic ligation via Sortase A, which covalently cross-linked substrate peptides present in each of the two hydrogel layers (**Figure 1d, Supplementary Movie 1**).

**Figure 1.**
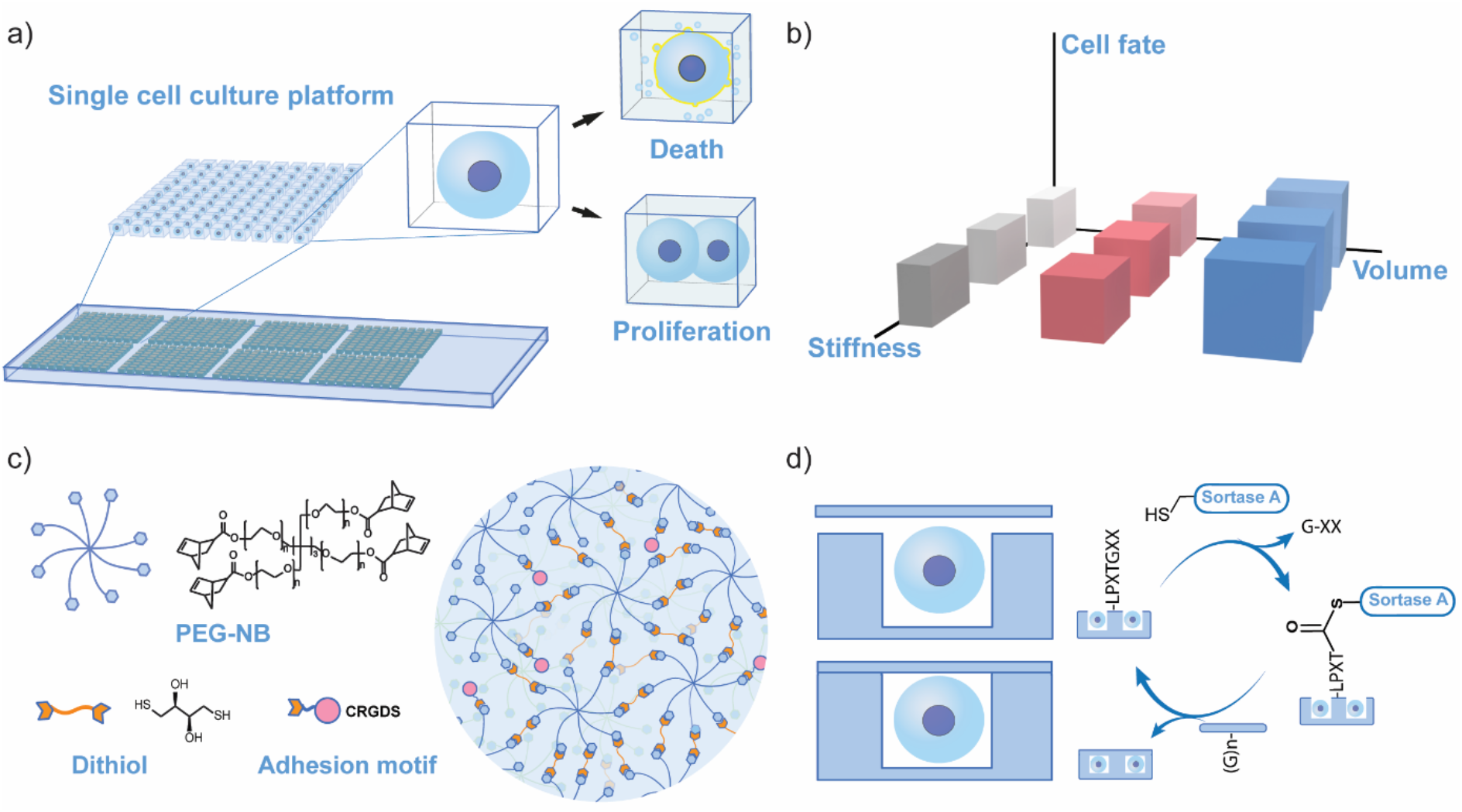
Design of the single cell culture platform with arrays of niches of defined volume and stiffness. a) The single cell platform was designed to fit on a standard glass slide. The hydrogel was patterned with structured arrays of geometrically defined micron-sized niches that can host single cells. We used this platform to investigate the role of geometric confinement and niche properties on hMSC fate, cell viability, and proliferation in 3D. b) We studied the specific effects of 3D niche volume and stiffness on hMSC viability, proliferation, and YAP localization (cell fate). We also compared these confined cells to unconfined hMSCs on 2D hydrogels of the same stiffness. c) The hydrogels were formed via thiol–ene photopolymerization between norbornene-functionalized star PEG macromers (PEG-NB) and a dithiol. The hydrogels were functionalized with CRGDS to facilitate cell adhesion. d) To create full 3D confinement of the encapsulated cells, the single cell niches were sealed using enzymatic ligation. The bacterial enzyme Sortase A cross-linked two substrate peptides present in the array base and a non-patterned lid. First, the sulfhydryl group of Sortase attacks the T-G amide linkage in Ac-GCRE-DDD-LPMTGG to form a LPMT-Sortase thioester intermediate, which is then attacked by the N-terminus of GGGG-LERCL to form a covalent amide cross-link between the two gel stabs, with the structure Ac-GC(PEG)RE-DDD-LPMT-GGGG-LERC(PEG)L. In this manner, the hydrogel lid was linked to the micropatterned array enclosing the niches.

We fabricated the microniche arrays using traditional microfabrication and soft lithography techniques (**Figure 2a**). The micropattern was designed in AutoCAD and generated on a silicon wafer via photolithography using SU-8 photoresist (**Figure 2b**). The silicon wafer was used as a master for the fabrication of intermediate polydimethylsiloxane (PDMS) molds. Teflon molds were cast from the intermediate PDMS patterns to avoid hydrogel patterning directly on PDMS. In our experience, the radical mediated thiol–ene polymerization was inhibited at the interface of PDMS, likely due to oxygen inhibition of primary radicals, limiting pattern transfer fidelity (**Figure S1**). We then cast microniche arrays in the PEG-based hydrogels using the Teflon molds and confirmed transfer by visualization of Rhodamine-labelled hydrogels via transmitted light and confocal microscopy (**Figure 2c,d**). The elastic modulus (and stiffness) of the base hydrogel was controlled by the polymer concentration in the precursor solution. Shear rheometry quantified the mechanical properties of the hydrogels in situ during formation and at equilibrium swelling. We selected hydrogel formulations with three distinct moduli at equilibrium swelling, *E* = 6, 16, or 30 kPa, referred to as soft, medium, and stiff gels (**Figure 2e**). These moduli were selected based on the stiffness range of physiologic hMSC microenvironments and prior in vitro investigations of hMSC mechanobiology.^31^ Previously, low moduli PEG-based substrates (*E* < 10 kPa) mechanically deactivated hMSCs (lower proliferation rates and cytoplasmic YAP localization) while stiff PEG-based substrates (*E* > 10 kPa) activated hMSCs (increased proliferation rates and nuclear YAP localization).^32^

**Figure 2.**
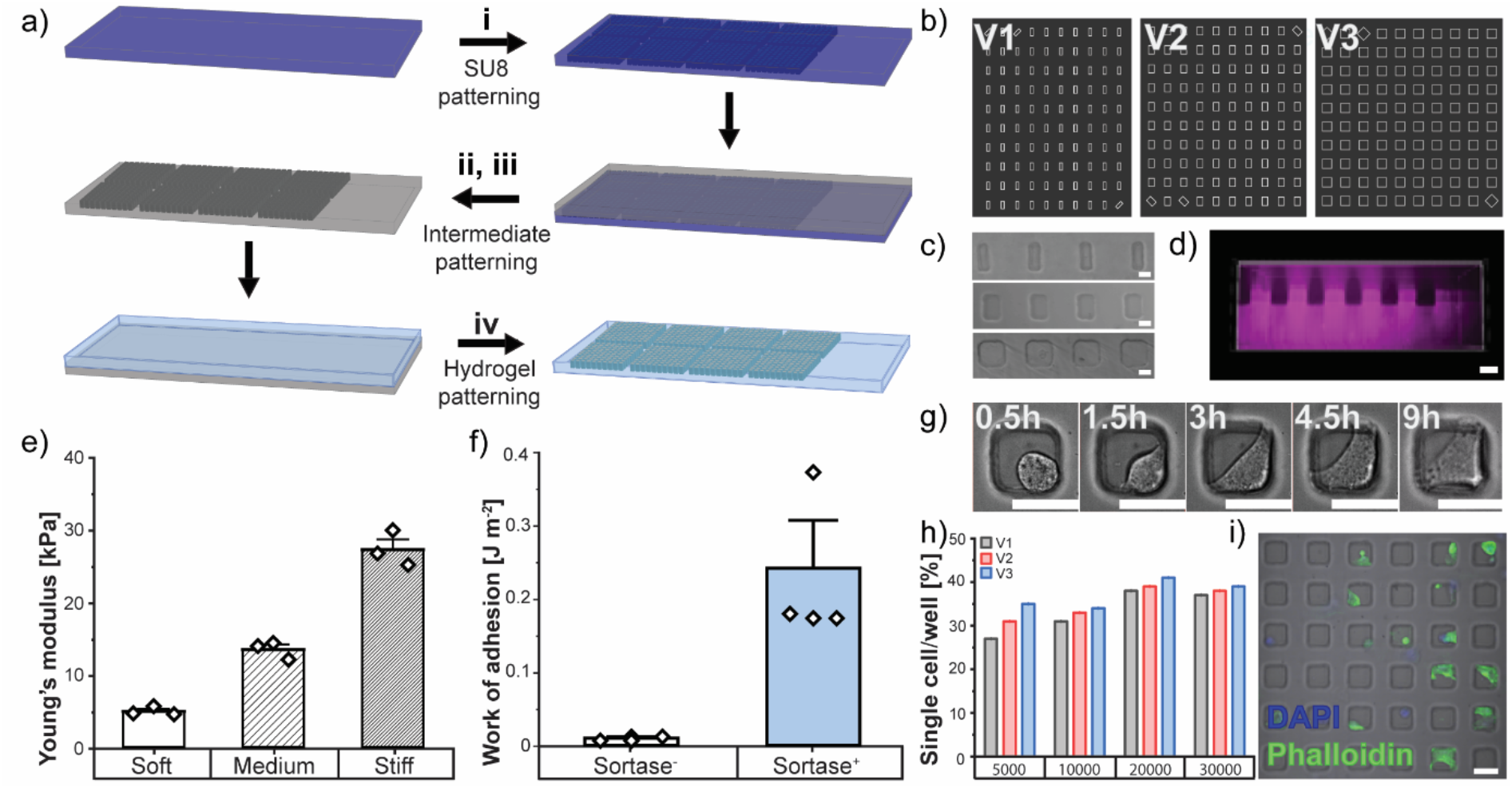
Fabrication of single cell microniche arrays and encapsulation of single hMSCs. a) The microfabrication process of the microniche arrays: i. Fabrication of the silicon master was performed via standard soft photolithography techniques. ii., iii. Fabrication of the intermediate PDMS mold from the silicon master was followed by the casting of the Teflon mold. iv. The Teflon mold was used to generate the patterned hydrogel. b) Images showing fragments of the AutoCad masks for V1 (20 x 50 x 35 μm^3^), V2 (35 x 50 x 35 μm^3^), and V3 (50 x 50 x 50 μm^3^) niches. The microwells were structured in 10×10 arrays to facilitate position identification and rapid image analysis. c) Transmitted light image of the resulting hydrogel pattern showing x–y shape fidelity between the original pattern and the generated microarray. Scale bar, 20 μm. d) Confocal image of a labelled hydrogel demonstrating shape fidelity in the z-dimension. The hydrogel was visualized through the incorporation of a thiolated Rhodamine dye that was covalently attached to the PEG backbone. Scale bar, 50 μm. e) The Young’s modulus of the hydrogel networks was controlled by the polymer fraction yielding soft (3.5 wt%; *E* = 6 kPa), medium (5 wt%; *E* = 16 kPa) and stiff (6 wt%; *E* = 30 kPa) gels. f) The lids adhered to the microarrays following Sortase A linking. The work of adhesion between the two hydrogels was quantified using pull-off experiments with and without the application of the bacterial enzyme Sortase A. The adhesion was sufficient for gels to remain adhered over the time course of the experiment. g) Live imaging showing a representative hMSC spreading within a V3 niche. The cell spread and stabilized its shape over the course of 9 h. Scale bar, 50 μm. h) The highest single cell occupation in the microniches was observed for 20 000 cells cm^-2^. i) Cells seeded in V3 niches after 3 days of culture. The actin cytoskeleton was labelled with Phalloidin-AF488 and the nucleus with DAPI. Scale bar, 50 μm.

The microniche arrays were sealed with a hydrogel made with the same formulation and material properties as the niche environment. To adhere the non-patterned hydrogel layer to the microniche base, the sealing hydrogel layer was modified with one substrate for Sortase A (GGGG-LERCL-NH_2_; 688 μg mL^-1^, 800 μM) and the micropatterned hydrogel was modified a complementary substrate (Ac-GCRE-DDD-LPMTGG-NH_2_; 1125 μg mL^-1^, 800 μM). The work of adhesion between gels was assessed using pull-off tests measuring stable adhesion between hydrogel surfaces in presence of Sortase A enzyme (**Figure 2f**). The Sortase-mediated bonding adhered the sealing hydrogel to the base throughout the time course of cell culture and in a manner that did not alter the network structure of the gels at the cell–material interface or interfere with cell function.

To investigate the effect of the degree of confinement on cell function, we generated microarray patterns with three distinct niche volumes: 35, 61, and 125 x 10^3^ μm^3^, further denoted as V1, V2, and V3 niches. The dimensions of the single cell niches were large enough to accommodate both the nuclear and cytoplasmic volumes of rounded hMSCs, with the smallest niche dimension (20 μm) exceeding the mean radius of hMSC nuclei (14.1 ± 2.0 μm; **Figure S2**). The niches were designed in this manner so that the cells and their nuclei would not be deformed upon encapsulation. Single cells were able to spread and fill smaller niches, while having sufficient space to spread, grow, and proliferate in larger niches. However, the largest niche dimension was smaller than the mean hMSC spread length in 2D culture (>50 μm, soft gels; >100 μm, stiff gels) in order to physically confine cells within the 3D microenvironments (**Figure S3**).^33^ In 3D, the maximum cell spread length was restricted to the longest dimension of the niche (~64 μm, V1; ~70 μm, V2; ~86 μm, V3).

After preparing the microarrays, we seeded cells into the niches by applying a suspension of hMSCs on the surface of the patterned hydrogel. To determine a suitable seeding density for single cell occupation, we seeded the cells in the niches at different concentrations (5 000–30 000 cells mL^-1^, 1 mL cm^-2^). We allowed the cells to sediment into the microwells under a constant gentle shaking on an orbital shaker (60–70 rpm) for 15 min. Once the cells settled into the microwells, the platform was rinsed with culture medium to remove excess cells from the surface. The niches were then sealed with the non-patterned hydrogel layer. Cell spreading was observed in the niches within 3 h after seeding (**Figure 2g, S4, Supplementary Movie 2**). We found that seeding 20 000 cells cm^-2^ of patterned surface resulted in the highest rate of single cell occupancy (~40%) for all niche geometries (**Figure 2h**). On day 3, the majority of the cells had spread within the niches attaining 3D morphologies (**Figure 2i, S5, Supplementary Movie 3**). In the V1 niches, spread cells often occupied the full microniche, whereas cells in V3 niches only spread to occupy a fraction of the niche (**Figure S6**).

### Geometric confinement and matrix stiffness affect hMSC viability

Having established the microarray platform for single hMSC culture, we investigated the effects of geometric confinement and matrix stiffness on hMSC fate. Initially, we employed our platform to investigate how 3D confinement and niche stiffness affect cell viability. We monitored cell viability with ethidium homodimer (EthD-1) staining, a DNA intercalator that cannot cross the membrane of viable cells. hMSCs that did not stain for EthD-1 (EthD-1^-^) were counted as viable and hMSCs that stained for EthD-1 (EthD-1^+^) were counted as dead. We screened viability across all niche volumes (V1, V2, and V3) in low, medium, and high stiffness gels. On day 1, hMSCs cultured in the 3D microniches exhibited a mean viability ≥80% for all conditions (**Figure 3a**). On day 3, hMSC viability varied with microniche volume and matrix stiffness. The number of viable cells decreased in low and medium stiffness niches (72.3 ± 13.6% and 79.5 ± 6.8%, respectively) and remained high in high stiffness niches (93.6 ± 5.5%). A weak and non-significant trend in viability was observed for niche volume. On day 3, the V1 niches at low stiffness exhibited heterogeneous results with decreased viability (59.1 ± 18.9%). Whereas the V3 niches with high stiffness exhibited the highest viability observed in 3D (97.5 ± 2.1%; **Figure 3b**). These data suggested that cells respond to 3D confinement, that niche volume and stiffness influence confinement sensing, and that these effects may be related to mechanical signaling.

**Figure 3.**
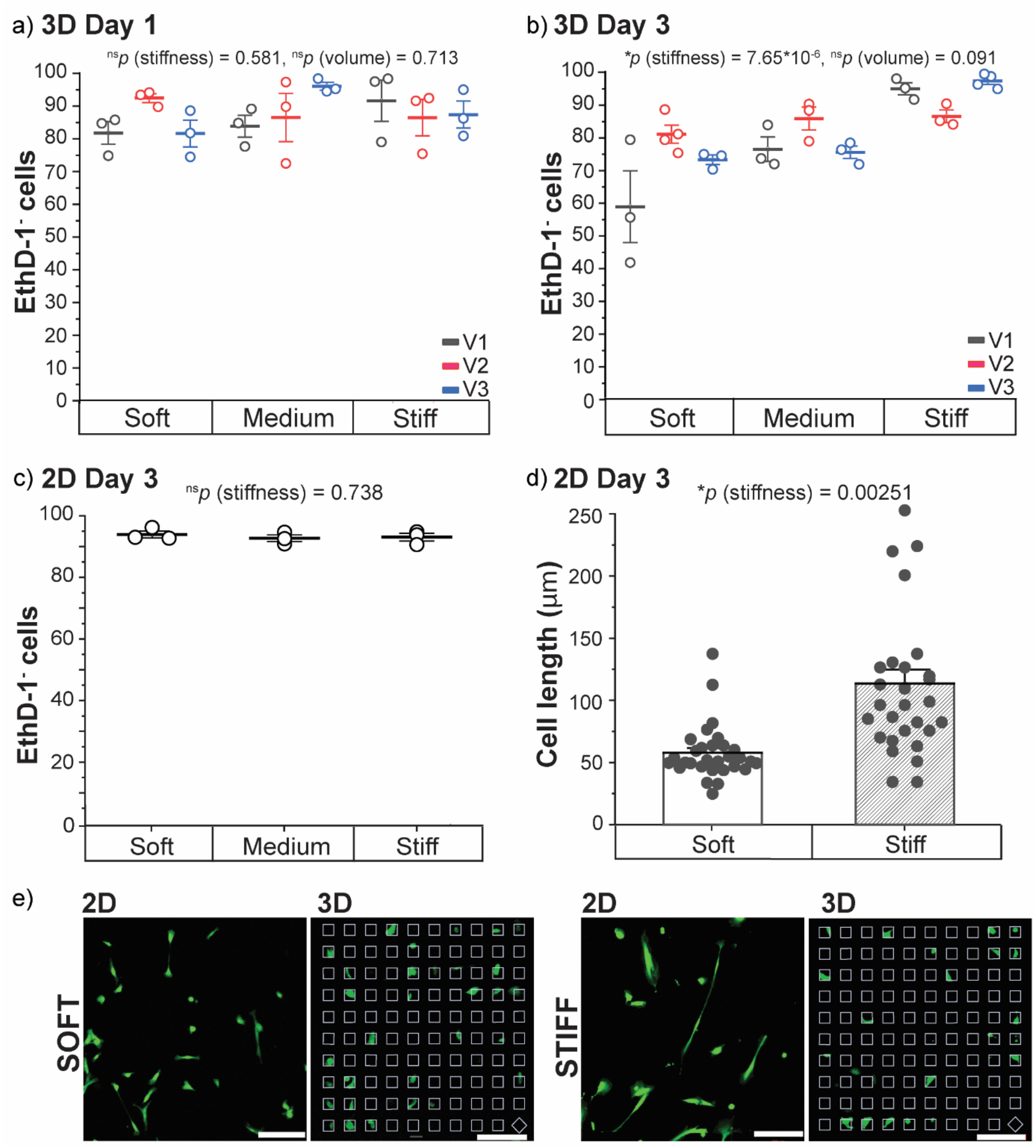
3D confinement decreased cell viability. a) hMSCs were viable (>80%) in all conditions in 3D microniche culture after 1 day (n = 3; at least 500 cells per condition; comparisons between stiffness and size groups were made using two-way ANOVA with Tukey’s test for post-hoc analysis). b) After 3 days, the high stiffness niches displayed the highest numbers of viable (EthD-1^-^) cells (n = 3; at least 500 cells per condition; comparisons between stiffness and size groups were made using two-way ANOVA with Tukey’s test for post-hoc analysis). c) On 2D gels, cell viability was close to 100% for all conditions after 3 days in culture (n = 3; at least 2000 cells per condition; comparisons between stiffness groups were made using oneway ANOVA, with Tukey’s test for post-hoc analysis). d) In unconfined 2D culture, hMSCs were more elongated on stiff gels as compared with soft gels (n = 3; 30 cells per condition; comparison of means between groups was made using a Two-Sample t-Test). e) In both conditions, hMSC spreading was confined in the 3D microniches relative to unconfined 2D gels. In 2D, cell spreading increased with gel stiffness and, in all conditions, cells adopted spread morphologies with mean cell lengths greater than the maximum available distance in the V1, V2, and V3 niches. In 3D, cell spreading was constrained by the geometry of the microniche, providing geometric confinement of the cells. Representative images of hMSC morphology on day 3 in 2D and 3D culture. Cell cytoplasm was imaged with Calcein AM. Scale bars, 200 μm. For a-c) plots represent mean ± s.e.m., for d) bar plots represent mean + s.e.m.

To further characterize the effect of geometric confinement on cell viability, we compared the viability of hMSCs in 3D microniches (confined) with hMSCs on 2D gels (unconfined) of the same stiffness. Cells exhibited high viability (≥ 90%) in all cases on 2D gels (**Figure 3c**). hMSC morphology varied with substrate stiffness; cells were less spread and more rounded on low stiffness gels and more spread on high stiffness gels with an average spread length of 57.9 ± 4.1 and 114.1 ± 11.2 μm, respectively (**Figure 3d**). However, the most notable effect was the overall increase in hMSC viability in 2D as compared with hMSCs in 3D microniches. While the specific niche volume did not have a significant effect on cell viability, 3D confinement dramatically reduced the overall viability of hMSCs. This could have been caused, in part, by mechanical effects during the assembly of the microarrays; however, as the majority of cells exhibited a decrease in viability after 3 days of culture in niches, we hypothesized that the cells actively sensed the confinement and that mechanosensitive signaling related this signal to coordinated cell fate decisions.

### hMSC proliferation decreases upon 3D confinement

Having shown that the rates of cell death increased upon confinement, we then investigated the proliferative capacity of hMSCs in the confined microniches as a function of niche stiffness and volume. Seminal results demonstrated that cell proliferation rates decreased with decreasing cell adhesive area on 2D adhesive islands (75–3 000 μm^2^).^34^ Other findings demonstrated that proliferation in 3D—porous ECM-derived and synthetic bulk hydrogels—is reduced with respect to 2D culture.^35,36^ Therefore, we hypothesized that proliferation rates would decrease upon confinement in 3D niches relative to 2D and that the rate of proliferation would correlate positively with niche volume. Further, we hypothesized that increased matrix stiffness would promote hMSC proliferation similar to observations in vivo or in 2D culture.^37^ Increased tissue stiffness has been associated with increased cell proliferation and invasion in different types of cancers.^38,39^ Similarly, stiffness has been shown to activate proliferation in vitro on 2D substrates and in porous ECM-derived hydrogels in 3D.^37,40^ However, conflicting effects have been observed in bulk 3D biomaterials showing a decrease in cell proliferation with increasing stiffness.^35^ This may be caused by the inherent increase in cell confinement at high stiffnesses in traditional bulk hydrogels.

To further study how confinement affects proliferation, we cultured hMSCs in our 3D microniches for 1 and 3 days and assessed proliferation by calculating the fraction of cells in S-phase via 5-ethynyl-2’-deoxyuridine (EdU) staining (**Figure S7**). Strikingly, the mean percentage of proliferating cells on day 1 in all 3D conditions was below 10%, which is much lower than hMSC proliferation observed in standard 2D culture (57.0 ± 3.0%; P7 hMSCs on TCPS) (**Figure 4a**).^41^ The mean value of EdU^+^ hMSCs in the 3D niches varied from 4.6 ± 2.3% to 8.1 ± 1.4%. At this early timepoint, the percent of proliferating cells increased with niche volume but did not exhibit an obvious trend with niche stiffness. On day 3, the proliferation rate settled below 10% and was lowest for the smallest (V1) niches (**Figure 4b**). The percent of EdU^+^ cells remained at ~7–8% in the V3 niches but dropped to 3.6 ± 0.4% in the V1 low stiffness niches. Overall, hMSCs exhibited the highest proliferation rates in V2 and V3 niches at medium and high stiffnesses (7.7 ± 0.9% and 7.9 ± 0.7% in high stiffness niches, respectively). Niche stiffness had a significant effect on proliferation rates resulting in the lowest number of EdU^+^ cells in low stiffness arrays, and proliferation rates increased with stiffness for all size niches (**Table S13–15**). The effect of stiffness was most pronounced in V1 and V2 niches.

**Figure 4.**
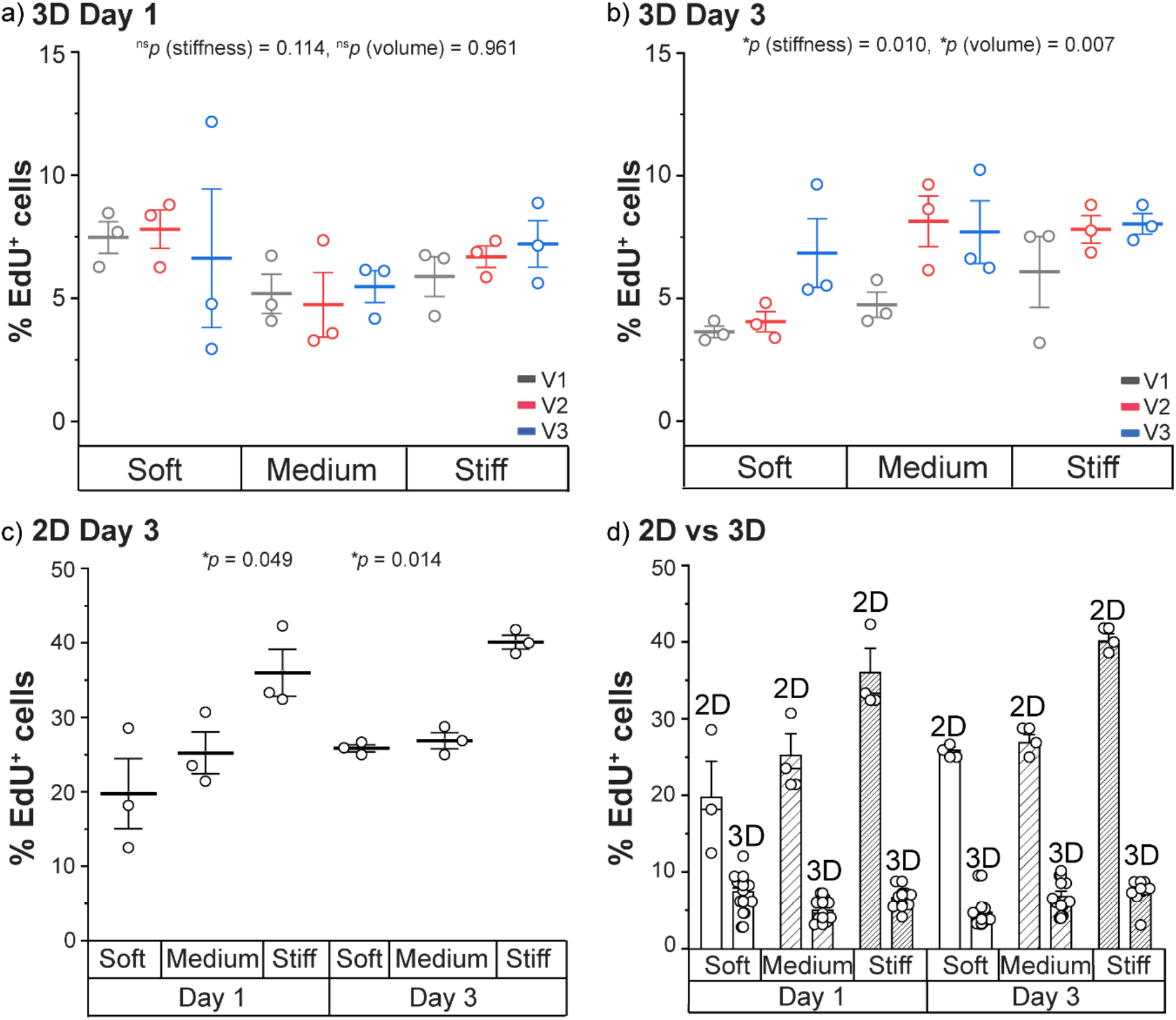
3D confinement downregulated hMSC proliferation. hMSCs were cultured on 2D substrates (unconfined) and in 3D microenvironments (confined) for 1 and 3 days. a) In 3D, cells displayed mean proliferation rates below 10% after 1 day in all niche conditions. (n = 3; at least 500 cells per condition; comparisons between stiffness and volume groups were made using a two-way ANOVA with Tukey’s test for post-hoc analysis). b) After 3 days, proliferation in the 3D niches remained low, further decreasing for V1 niches. (n = 3; at least 500 cells per condition; comparisons between stiffness and volume groups were made using a two-way ANOVA with Tukey’s test for post-hoc analysis). c) On day 1 in 2D culture, the number of EdU^+^ cells varied from ~20 to 36%; the proliferation rate increased with substrate stiffness. After 3 days, 2D proliferation increased further to above 25% for soft and 40% for stiff gels (n = 3; comparisons between stiffness groups were made using a one-way ANOVA with Tukey’s test for post-hoc analysis). d) After 1 and 3 days of culture, hMSC proliferation rates in the confined 3D microniches were substantially than values for unconfined hMSCs in 2D culture. For a-d) plots represent mean ± s.e.m. For e) bar plots represent mean + s.e.m.

To relate these observations to geometric confinement, we compared proliferation values for hMSCs in 3D microniches with unconfined hMSCs on 2D gels of the same mechanical properties. After 24 h in culture, the percent of EdU^+^ cells on 2D substrates varied from 19.8 ± 8.1% to 36.0 ± 5.5%; proliferation rates increased with substrate stiffness (**Figure 4c**). On day 3, 2D proliferation increased further to 25.9 ± 0.8% for soft and 40.1 ± 1.6% for stiff gels (**Figure 4c**). Overall proliferation rates were markedly higher for unconfined hMSCs on 2D gels than for confined hMSCs in 3D microniches for all stiffnesses tested (**Figure 4d**). These results indicated that 3D geometric confinement and matrix stiffness affect stem cell life and death.

### Geometric confinement and niche stiffness regulate YAP localization in 3D

Our observations demonstrated that stem cell fate was influenced by geometric confinement in 3D—proliferation rates decreased upon mechanical confinement and rates of cell death increased, and both of these effects were coupled to niche volume and stiffness. Since the cells on 2D displayed average spread lengths that exceeded the longest dimension of the niches, we hypothesized that physical restriction in confined microniches may result in decreased mechanotransduction by inhibiting cell elongation and cytoskeleton maturation. Of the many effectors involved in mechanosensing, we opted to investigate YAP, which shares homology with transcriptional coactivator with PDZ-binding motif (TAZ), as a potential mediator of the response to 3D confinement. YAP/TAZ nuclear activity is governed, in part, by mechanical cues that are transduced through the cytoskeleton and YAP/TAZ localization depends on internal cell tension.^42,43^ Previously, YAP/TAZ activation has been shown to depend on the available adhesive area on confined 2D micropatterned fibronectin islands; YAP/TAZ was activated in cells on large islands and inactivated on smaller islands.^44^

To investigate if the effects of mechanical confinement on cell fate were indeed mediated by YAP signaling, we characterized YAP localization (nuclear or cytoplasmic) in hMSCs cultured in confined 3D microniches and on unconfined 2D gels. The nucleo-cytoplasmic distribution of YAP is not a binary state but rather a time capture of continuous nuclear and cytoplasmic shuttling dynamics.^45^ We quantified YAP distribution by calculating nuclear and cytoplasmic mean fluorescent intensities, further defined as the nuclear/cytoplasmic ratio (N/C); we defined YAP nuclear localization at N/C values above 1.7 (**Figure 5a, S8**). YAP localization was assessed after 1 and 3 days in culture. First, we quantified YAP localization for hMSCs in 2D. As expected, the N/C scaled with the stiffness of the 2D substrates (**Figure 5b**). On day 3, the mean N/C ratio on soft gels was low (1.2 ± 0.5) indicating mechanical deactivation. Increased mean N/C values were observed on medium and stiff gels (1.8 ± 0.8 for medium and 1.7 ± 0.7 for stiff niches).

**Figure 5.**
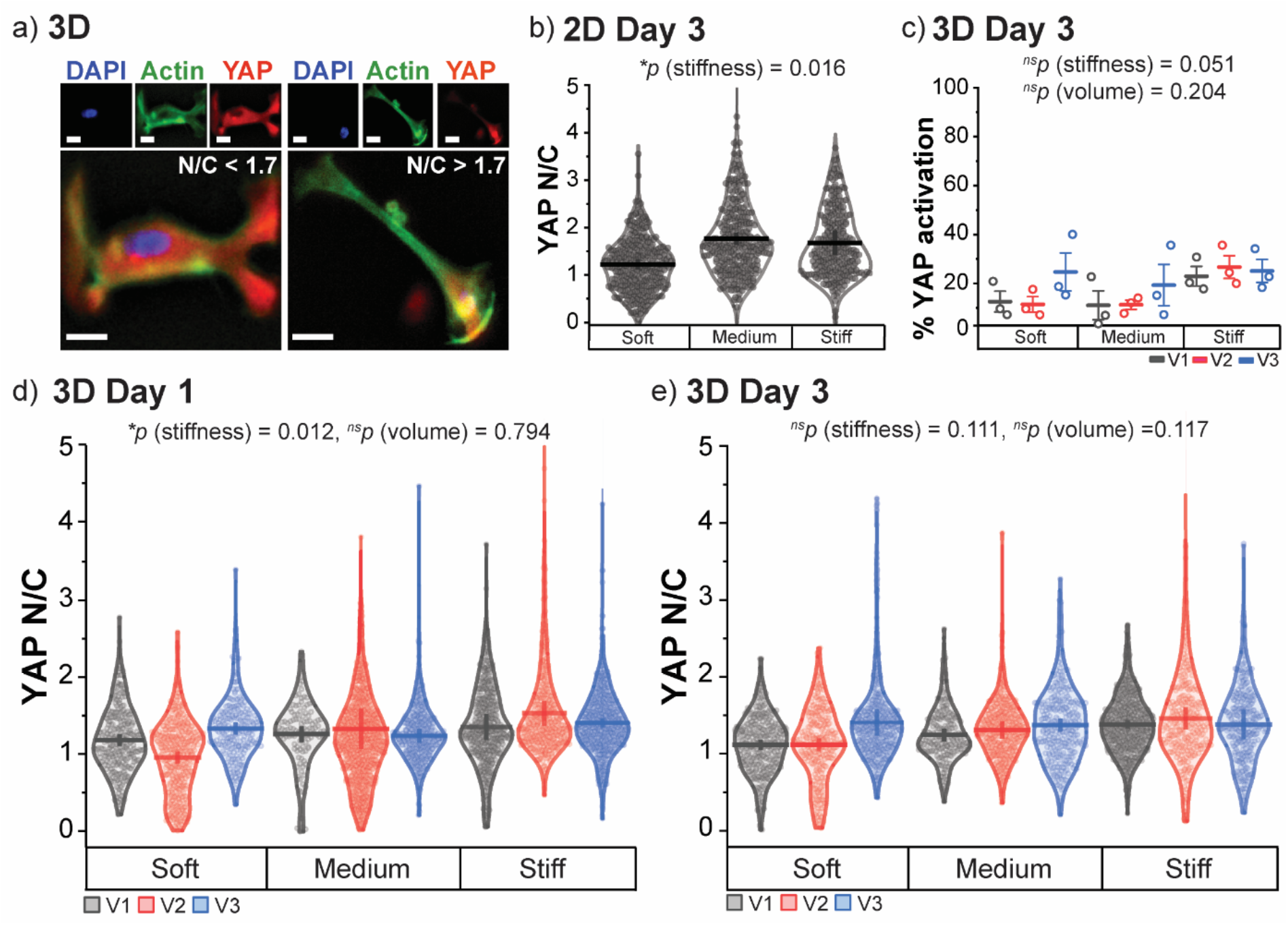
hMSCs were mechanically deactivated upon 3D confinement. a) Representative images of hMSCs in confined 3D V3 microniches with cytoplasmic (left; N/C ratio < 1.7) and nuclear (right; N/C ratio > 1.7) localization of YAP. Scale bars, 10 μm. b) During unconfined 2D culture on day 3, YAP N/C ratio (YAP/TAZ localization) increased gradually from ~1.3 for soft gels to ~1.7 for medium and stiff gels (n = 3; at least 250 cells per condition, comparisons between stiffness groups were made using a one-way ANOVA with Tukey’s test for post-hoc analysis). c) In confined 3D microniches on day 3, mean YAP activation varied from ~8% in soft V2 niches to ~24% in stiff V2 niches. Data represent mean ± s.e.m. (n = 3; at least 250 cells per condition; comparisons between stiffness and size groups were made using a two-way ANOVA with Tukey’s test for post-hoc analysis). d) The YAP N/C ratio of hMSCs after 1 day in 3D culture varied with niche volume and stiffness. All values were suppressed relative to similar conditions for unconfined hMSCs in 2D culture. e) After 3 days in 3D microniches, the YAP N/C ratio of hMSCs increased in larger volume niches for soft and medium gels. The YAP N/C ratio was elevated in stiff niches, independent of niche volume. However, the YAP N/C ratios remained suppressed relative to hMSCs on unconfined 2D gels with the same stiffness. For d) and e) the plots represent Kernel Smooth (KS) distributions with mean and 95% confidence interval of the mean (n = 3; at least 100 (d) or 250 (e) cells per condition, comparisons between stiffness and size groups were made using two-way ANOVA with Tukey’s test for post-hoc analysis).

In confined 3D microniches, hMSCs displayed mean YAP N/C ratios below the activation limit of 1.7 for all niche conditions on days 1 and 3. This suggested a general mechanical deactivation for confined cells as compared with unconfined cells in 2D culture. The mean percentage of YAP activation varied from 8.2 ± 5.6% in soft V2 niches to 23.9 ± 8.3% for stiff V3 niches (**Figure 5c**). On day 1, mean YAP N/C ratios did not correlate with niche volume and exhibited a weak trend with stiffness (**Figure 5d**). On day 3, the cells showed increased nuclear YAP localization in larger niches for soft and medium gels (**Figure 5e**). Interestingly, stiffness had a dominant effect on YAP N/C ratio: activation and N/C ratios were highest in stiff niches compared with other 3D microniche conditions, and the N/C ratio was independent of niche volume at this stiffness. Overall, suppressed YAP activation (mechanical deactivation, decreased YAP N/C ratios) in 3D niches correlated with our viability and proliferation measurements. It has been shown previously that proliferation correlates with mechanical activation of cells, which can be regulated through YAP signaling.^46^ Furthermore, YAP activation is a known suppressor of cell death.^47,48^ This indicates, that the changes in cell function within confined 3D environments may be in part regulated through YAP-mediated mechanotransduction.

## Discussion

When culturing cells outside of the body, the context matters and functional differences are observed between 2D and 3D culture.^49,50^ In 2D, cell death rates are generally low and proliferation rates are often supraphysiologic.^51,52^ In 3D, proliferation rates often decrease and cell death rates increase.^35,36^ These differences in cell function are often attributed to changes in cell polarization, adhesion, and uptake of soluble factors.^2^ Recently, an additional focus has been placed on physical confinement of cells in 3D as a factor that contributes to the context-dependent differences in cell fate.^26^ Seminal work on micropatterned 2D substrates demonstrated the role of confinement in directing cell fate and function.^7,9,53,54^ Further studies corroborated the link between cell confinement and fate suggesting that these effects are, in part, regulated by mechanotransduction.^7,9,53^ Studying the effects of confinement on cell fate in 3D is more challenging due to the general lack of control over geometric parameters of the cell microenvironment.

As an alternative, micropatterned 3D platforms enable the culture of individual cells in defined microwells.^55^ Here, we developed a platform to culture and monitor single hMSCs in confined microenvironments in order to investigate systematically the effect of geometric confinement on stem cell fate in 3D. We studied cell fate in unconfined 2D culture and confined 3D culture across a range of niche volumes and stiffnesses. In 3D, cell death increased with confinement and was significantly elevated relative to 2D culture. In addition, we observed a marked decrease in cell proliferation in 3D microniches as compared with 2D culture across all niche volumes and stiffnesses. Even though the largest volume niches exceeded the average hMSC volume, proliferation in 3D was significantly lower than in 2D. The observed proliferation rates in 3D were similar to in vivo proliferation rates in soft tissue, which typically do not exceed 10% of S-phase cells except during tissue development, regeneration, or tumor growth.^56–58^ These data indicate that cells sensed confinement in their microenvironment even when the niche volume exceeded that of an individual cell and that confinement affected the rates of cell life and death.

We hypothesized that the observed effects of physical confinement may be mediated by mechanosensitive signaling pathways.^53,59^ We investigated YAP localization as a function of confinement and stiffness. YAP activity is, in part, governed by mechanical loading transduced through cytoskeleton organization, and reduced YAP activation in hMSCs has correlated with reduced viability and proliferation in 2D culture.^60–62^ Consistent with our proliferation results, YAP activation in 3D was reduced relative to 2D, indicating that cells experienced lower mechanical stresses upon confinement. Traction dynamics of actin stress fibers are related to their length and in 2D cells balance external forces by adjusting the stiffness and organization of cytoskeleton, including its length, to substrate stiffness.^40,63,64^ 2D confinement of cells to smaller spread areas has been shown to suppress stress fiber formation.^59,65^ 3D niches impose similar physical restrictions on cell elongation, length, and organization of actin stress fibers. Therefore, 3D confinement may act similarly to 2D confinement hindering actin stress fiber formation resulting in dampened cell contractility and stiffness sensing.^66,55^ Correspondingly, both proliferation and YAP activation in 3D niches scaled positively with niche volume—larger niches allowed for increased cell elongation. Related studies that assessed proliferation with respect to the diameter of scaffold porosity have shown similar results—larger pore sizes stimulated cell proliferation.^67,68^

Furthermore, in the microwells, hMSCs spread in 3D, which results in less polarized cell spreading as compared with 2D; cell polarity has been shown to alter mechanotransduction.^69,70^ Namely, cell polarization on 2D substrates has been shown to induce stretching of the nuclear envelope.^32^ Nuclear flattening increases nuclear pore diameter on the cytosol side and reduces it on the nucleoplasmic side, lowering the export rate of YAP/TAZ in comparison to the import rate, favoring nuclear accumulation.^71^ Chaudhuri and coworkers have shown that, in contrast to 2D substrates, 3D conditions allow for non-polarized adhesion, imposing lower stretching forces on the nucleus and resulting in lower nuclear cross-sectional area.^72^ In our niches, hMSCs spread in 3D given the isotropic distribution of adhesive cues. We hypothesize that limited cell elongation and 3D morphology of cells in our confined niches may correspond to a less polarized nuclear conformation, reducing YAP/TAZ activation at high stiffness as compared with 2D culture.

Lastly, increased cell death was most apparent in soft and medium stiffness niches, while, in stiff niches, cells maintained high viability close to 2D levels. Across all conditions, proliferation increased with niche stiffness. In 2D, increased stiffness stimulates cell survival and proliferation due to elevated cell tension transduced through the cytoskeleton and contractile machinery of the cell.^51,73^ Although stiffness sensing in 3D niches might be reduced due to inhibited spreading, mechanotransduction is also regulated by the size and stability of focal adhesions, myosin contractility, and abundance of stress fibers, which are typically downregulated on low stiffness substrates.^74^ Thus, mechanotransduction may have been upregulated in stiff niches irrespective of cell elongation, resulting in the relative increase in cell viability and proliferation.

## Conclusion

In conclusion, this study improves our understanding of how the physical milieu regulates stem cell function and fate. By culturing hMSCs in microengineered niches, we demonstrated the role of geometric confinement and niche stiffness on hMSC proliferation and viability in 3D. Importantly, we observed significantly increased cell death rates and decreased rates of proliferation in hMSCs cultured within confined 3D microniches as compared with cells on unconfined 2D gels. We related the observed effects of geometric confinement and niche stiffness on cell fate to YAP localization, indicating that mechanotransduction pathways mediate these effects. Further investigation is needed in order to quantitatively describe the effects of confinement on the mechanical activation in cells and how it relates to cytoskeletal organization, focal adhesion formation, and generation of traction forces. A general understanding of how cell behavior is affected by geometric confinement will be an important step toward mapping a complete regulation profile of in vivo cell fate. While we focused here on the role of confinement on single cell fate, 3D confinement is likely to be involved also in the regulation of multicellular structures influencing growth, development, and homeostatic profiles of tissues.

## Methods

### Hydrogel platform design

#### Synthesis of norbornene-functionalized PEG

8-arm PEG amine (M_n_ ~10 kDa; 4 g, 0.4 mmol PEG, 3.2 mmol NH_2_, 1 eq. NH_2_; JenKem USA) was dissolved in anhydrous dimethylformamide (DMF; 5 mL; Sigma-Aldrich) and purged with argon. N,N-Diisopropylethylamine (DIPEA; 2.23 mL, 12.8 mmol, 4 eq.; Sigma Aldrich) was added to the PEG solution followed by the addition of 1-[bis(dimethylamino)methylene]-1H-1,2,3-triazolo[4,5-b]pyridinium 3-oxide hexafluorophosphate] (HATU; 2.43 g, 6.4 mmol, 2 eq.; Sigma-Aldrich). Next, 5-norbornene-2-carboxylic acid (1.56 ml, 12.8 mmol, 4 eq.; Sigma-Aldrich) was added to the solution and the reaction was stirred overnight at room temperature (RT) under argon atmosphere. The product was precipitated twice in diethyl ether (4 °C) and the precipitated polymer product was recovered and dialyzed in a dialysis membrane (MWCO 1000 g mol^-1^; Spectrum Laboratories) against dH_2_O for 3 days. The aqueous polymer solution was lyophilized obtaining the product in the form of a white amorphous solid. Functionalization of the 8-arm PEG with norbornene was determined to be above 95% via ^1^H NMR in CD_2_Cl_2_ by comparing the integrated areas under the peaks of the norbornene vinyl protons (δ = 6.0–6.3, m, 2H) and the PEG ether protons (δ = 3.5–3.9, m, 96H; **Figure S9**).

#### Sortase A production

The Sortase A pentamutant (eSrtA) in pET29 plasmid was kindly gifted by Prof. David Liu (Addgene plasmid # 75144). Production was performed in BL21 (DE3) *E. coli* (Invitrogen). We first inoculated 12 ml of lysogeny broth (LB; ThermoFisher) containing 50 μg/ml kanamycin (ThermoFisher). The inoculate was then incubated at 37°C, 180 rpm overnight. We then transferred 5 ml of this inoculate to 500 ml of LB + kanamycin for culture at 37°C, 180 rpm until OD600 = 0.8 (around 3h30). After induction with 0.2 mM Isopropyl-β-D-thiogalactoside (IPTG, Sigma-Aldrich), we further incubated at 16°C for 22 h, and cells were centrifuged and lysed with BugBuster (Merck), followed by purification on a His-Trap HP affinity column (GE Healthcare) using a gradient from 10 to 250 mM imidazole over 30 min in Tris buffered saline pH 7.5 in the presence of 1 mM betamercaptoethanol (Sigma-Aldrich). After concentration by centrifugation on Vivaspin (10000 g mol^-1^ MWCO; Sartorius), we further purified to remove endotoxins with Pierce high capacity endotoxin removal spin columns (ThermoFisher). The buffer was exchanged to TBS + 1 mM beta-mercaptoethanol (Sigma-Aldrich) by dialysis at 4 °C (1000 g mol^-1^ MWCO; Spectrum Laboratories), 4 times 6 h. Finally, TBS+20% glycerol (Sigma-Aldrich) was added dropwise over stirring to the protein solution until reaching 10% glycerol content, and after quantitation using area under the curve on analytical gel permeation chromatography (Agilent Aquagel 20 column, 7.8 × 300 mm, with 5 μm particle size, using TBS 0.5 ml/min as the eluent), the solution was further diluted with TBS+10% glycerol to obtain a 400 μM stock. This stock was sterilized by filtration at 0.2 μm, aliquoted, and stored at −80 °C.

#### Synthesis of Sortase A peptide substrates

The Sortase A substrate peptides Ac-GCRE-DDD-LPMTGG-NH_2_ (Sortase A Threonine-donor or SAT) and GGGG-LERCL-NH_2_ (Sortase A Glycine-donor or SAG) were synthesized on a Prelude X peptide synthesizer (GYROS Protein Technologies). Ac- and -NH_2_ refer to acetylated and amidated N- and C-termini respectively, the GCRE cassette provides a reactive cysteine for Michael addition in a hydrophilic block, DDD acts as a hydrophilic spacer, and LPMTGG is a highly reactive Sortase A substrate sequence. The alternative LERCL cysteine makes the peptide slightly less hydrophilic to facilitate purification (using a conventional cassette with G instead of L results in a peptide that elutes with the injection peak here). The solid phase supported synthesis used fluorenylmethoxycarbonyl (fmoc) protected amino acid building blocks, induction heating, and mixing was performed with nitrogen bubbling in all reaction steps. For a 0.5 mmol synthesis, 25 ml glass reaction vials and 12.5 ml of solvent/solution. High load Rink Amide MBHA Resin (GYROS Protein Technologies cat. RAM-100-HL) was swelled for 60 min in DMF at RT. Deprotection was performed with 20% piperidine in DMF treatment twice for 1 min at 50°C, followed by three DMF washes. Coupling was performed twice (thrice for first coupling) for 5 min at 50°C, using a reaction mix in DMF containing protected amino-acids 0.2 M stocks, HBTU 0.4 M stock, and N-methylmorpholine 0.4 M stock, in 40/20/40 proportions, for a total of 1 mmol (2 equivalents) amino acid per coupling. The resin was then washed three times with DMF before repeating the deprotection/coupling cycle. For N-terminus acetylation, the resin was treated *in situ* with 20% acetic anhydride in DMF for 10 min at RT. Finally, the resin was washed with dichloromethane (DCM; Sigma-Aldrich), dried with nitrogen for 60 min, and cleaved for 2h at RT. The cleaving solution for SAG (optimized to avoid side products from reaction with rink amide linker in addition to side reactions with cleaved protecting groups and cysteine oxidation) contained TFA/H2O/TIPS/EDDT in proportions 85/7.5/5/2.5. The cleaving solution for SAT (further optimized to avoid methionine oxidization) contained TFA/TIPS/EDDT/thioanisole/anisole in proportions 83.75/3.75/3.75/6.25/2.5, where TIPS (Sigma-Aldrich) is triisopropylsilane and EDDT is 2,2’-(ethylenedioxy)diethanethiol (Sigma-Aldrich). The crude peptides were the precipitated in ice cold diethyl ether (Et2O; Sigma-Aldrich), collected by centrifugation, further washed with ice cold Et2O, dried under nitrogen flow, and resuspended in a minimal amount of acetonitrile/water/TFA, followed by purification by preparative reverse-phase high-performance liquid chromatography (HPLC; Agilent 1260 infinity) on a 55 mm diameter C18-capped silica column (Agilent), using a gradient from 10 to 90% acetonitrile in water over 40 min in the presence of 0.1% TFA. LC-MS (high resolution, positive mode, **Figure S10**): SAG, calculated mass: 860 g mol^-1^; measured m/z: 860 [M]^+^, 1720 [2M+H]^+^; SAT, calculated mass: 1406.49 g mol^-1^; measured m/z: 704 [M+2H]^2+^, 1407 [M+H]^+^.

#### Synthesis of lithium phenyl-2,4,6-trimethylbenzoylphosphinate

Lithium phenyl-2,4,6-trimethylbenzoylphosphinate (LAP) was synthesized as described previously.^75^ 2,4,6-trimethylbenzoylchloride (3.2 g, 18 mmol) of was added dropwise to an equimolar amount of continuously stirred dimethyl phenylphosphonite (3.1 g; 18 mmol) at RT and under argon. The reaction was stirred overnight under argon atmosphere. Lithium bromide (6.1 g; 72 mmol) was dissolved in 100 mL of 2-butanone and added to the reaction mixture. The reaction was heated to 50 °C to induce product precipitation. After 10 min, a solid precipitate formed. The reaction was cooled to RT over 4 h and then filtered to recover the precipitate. The precipitate was washed 3 times with 2-butanone (50 mL) to remove unreacted lithium bromide and dried under vacuum. The product was recovered in near quantitative yield. ^1^H NMR in D_2_O: 7.57 (m, 2H), 7.42 (m, 1H), 7.33 (m, 2H), 6.74 (s, 2H), 2.09 (s, 3H), 1.88 (s, 6H) (**Figure S11**).

#### Rheological characterization

The cross-linking kinetics and mechanical properties of the PEG hydrogels (3–10 wt%) were quantified using a strain-controlled shear rheometer (MCR 502; Anton Paar). The hydrogel precursor solution was loaded between an 8 mm parallel plate geometry (PP-08) and a transparent bottom plate with a gap of 0.5 mm. The gel forming solution was cross-linked with upon exposure to collimated UV light (λ = 365 nm, I = 20 mW cm^-2^; M365L3-C1, ThorLabs). Storage (*G′*) and loss (*G”*) moduli were measured over time with oscillatory strain measurements at γ = 1% amplitude and with an angular frequency of ω = 1 Hz (**Figure S12**) within LVE. All measurements were carried out in the linear viscoelastic regime for the formed gels. The Young’s modulus (*E*) was estimated using the relation between shear and Young’s moduli for isotropic and homogeneous materials, *E* = 2*G′*(1+v) where v is the Poisson’s ratio for the material. For the PEG-based materials tested, v was assumed to be 0.5.^76^

Additionally, the rheological tests described above were repeated on hydrogels after equilibrium swelling (~24 h) in phosphate buffered saline (PBS) to determine the swollen *G′*. The measurements were performed on hydrogels which dimensions were matching 8 mm geometry (sandblasted surface).

#### Sigmacote coating

To coat a substrate with Sigmacote, the substrate (silicon master or glass slide) was first cleaned in strong soap (SDS, Sigma-Aldrich) and washed thoroughly with water, ethanol, and acetone. After air drying, the substrate was placed in a Petri dish (silicon mater) or in a glass staining container (glass slides) and submerged in Sigmacote (Sigma-Aldrich) for 5 min. The container was sealed with a lid to avoid evaporation of Sigmacote. Subsequently, Sigmacote was removed and the surface was washed again twice with dH_2_O and air dried. In addition, the substrate was dried in oven at 100 °C for 30 min to produce a durable coating. Used Sigmacote was stored in a glass container and reused.

#### Silanization with (3-mercaptopropyl)trimethoxysilane

3-(Trimethoxysilyl)propyl methacrylate (TMPMA; Sigma Aldrich) has been used to covalently link thiol–ene gels to glass slides. Hydrogels cast on methacrylated slides did not lift from the glass when immersed in medium during cell culture and remain attached during staining procedures. To prepare the glass slides for silanization, the glass slides were cleaned with SDS, rinsed thoroughly with water, washed with ethanol and acetone, and dried in an oven at 80 °C for an hour. 1 mL of TMPMA was diluted in 200 mL of ethanol and 6 mL of dilute acetic acid (1:10 glacial acetic acid:dH_2_O) was added to the mixture directly prior use. The glass slides were placed in a glass staining container and submerged in the activated TMPMA solution allowing full contact of the glass surface with the silane solution. The solution was allowed to react for 5 min. The excess solution was poured off and the glass slides were rinsed with ethanol to remove the residual reagent and dried in air at RT.

#### Hydrogel adhesion via enzymatic ligation

Two hydrogel slabs (0.1 and 0.5 mm) were prepared at the same polymer wt% including complementary Sortase A peptide substrates (SAG or SAT, 800 μM). Two glass slides were treated with Sigmacote (Sigma-Aldrich) and separated with a 0.1 or 0.5 mm silicone rubber spacer. The hydrogel precursor solution was injected between the glass slides and polymerized upon exposure to UV light (λ = 365 nm, I = 20 mW cm^-2^, *t* = 2 min) The surface of the hydrogel slab containing SAT peptide was dried in air and treated with Sortase A solution (20 μL, 86 mg mL^-1^, 4 μM). Directly after, the second hydrogel layer containing SAG peptide was placed on top of the thicker slab substrate and pressed gently. The adhered hydrogels were submerged in a minimal amount of DPBS buffer containing Ca^2+^ and stored in the incubator at 37 °C for half an hour. Adhesion was assessed visually upon mechanical agitation (**Supplementary Movie 1**).

The work of adhesion following enzymatic cross-linking of two PEG hydrogel layers was assessed using pull-off tests performed using the normal force transducer of the rheometer (MCR-502; Anton Paar). Pre-swollen gels formulated with SAG or SAT (as described above) were attached to the Peltier plate or the 8 mm parallel plate geometry (PP-08) of the rheometer using cyanoacrylate glue. The surface of the hydrogel attached to the Peltier plate was treated with 10 μM Sortase A solution in Dulbecco’s phosphate buffered saline (DPBS with Ca^2+^ and Mg^2+^; ThermoFisher). The gel surfaces were brought into contact and pressed together until a normal force of 0.8 N was registered. The temperature of the Peltier plate was adjusted to 37 °C and a ring of DPBS was applied around the gels. The protective hood was lowered, to mitigate solvent evaporation, and the gels were left to crosslink for 30 min. After that, the DPBS was removed, and the upper geometry was retracted (0.01 mm s^-1^) during which the normal force (F_n_) upon retraction was measured and F_n_–displacement curves were recorded (**Figure S13**). In order to calculate the work of adhesion (J m ^−2^) from the recorded curves, we integrated the retraction force as a function of the displacement, followed by dividing the resulting adhesion energy by the known contact area (surface area of geometry) at the interface.

### Microfabrication

#### Microfabrication of silicon master

The microniche arrays were designed in AutoCAD (Autodesk). The obtained designs were used to fabricate polyethylene terephthalate (PET) based transparency masks with soft photographic negative emulsion film, right reading emulsion up (JD-Photodata). The resulting mask was used to transfer the design to a silicon wafer via photolithography in a clean room facility. A layer of photoresist SU-8 50 (MicroChem Inc.) was spin-coated onto a plasma-cleaned silicon wafer. Manufacturer recommendations were followed for prebaking, lithography, postbaking, and photoresist development. The resulting silicon wafer contained a micropost pattern. The pattern profile was assessed using white light interferometry (WLI), confirming that the dimensions of the microposts matched the dimensions of the desired microniches [V1 (20 x 50 x 35 μm^3^), V2 (35 x 50 x 35 μm^3^), and V3 (50 x 50 x 50 μm^3^)]. The masters with correct dimensions were used for the fabrication of the intermediate molds.

#### PDMS mold fabrication

The silicon master fabricated in the previous step was used for casting intermediate PDMS molds. In short, Sylgard 184 (Dow Inc.) was mixed with Sylgard curing agent (Dow Inc.) at a 10:1 mixing ratio by weight. The mixture was placed under vacuum in a desiccator for 30 min to remove trapped air bubbles. The silicon master was placed in a Petri dish and the PDMS mixture was gently poured onto the silicon master avoiding bubble formation. The Petri dish containing both the master with PDMS prepolymer was placed in an oven at 70 °C and cured for 6 h. The generated PDMS with a microwell pattern was separated from the silicon wafer and used in the following steps.

#### Teflon mold fabrication

The PDMS mold obtained in the previous step was used to fabricate a Teflon mold. The patterned PDMS mold was placed in the glass Petri dish and the surface was covered with Teflon beads. The Petri dish with PDMS and Teflon beads was placed overnight in the oven at 200 °C. After cooling to RT and detachment of the two materials, the generated Teflon mold with microposts was released and used for hydrogel patterning to produce the microniche arrays.

#### Hydrogel patterning

The hydrogel precursor solution was prepared as described in the procedure for the preparation of 2D hydrogels. The silicon spacer (0.5 mm) was placed on the TMPMA treated glass slide and the Teflon mold was placed on top of the spacer. The hydrogel precursor solution was carefully injected between the glass slide and the Teflon mold until the whole interstitial volume was filled, avoiding the formation of air bubbles on the surface of the mold. The precursor solution was placed under UV light (λ = 365 nm, I = 20 mW cm^-2^) for 1 to 3 min depending on the hydrogel formulation. Teflon mold was carefully removed from the hydrogel using a spatula as a lever while minimizing shear forces on the hydrogel. The patterned hydrogel adhered to the TMPMA-treated glass slide was immersed in PBS until further use.

#### Hydrogel pattern assessment

The hydrogel profile was analyzed using fluorescence confocal microscopy. In order to visualize the hydrogel profile, the hydrogel was labeled with acryloxyethyl thiocarbamoyl Rhodamine B (Sigma-Aldrich; **Figure S1**).^77^ The acrylated Rhodamine B was dissolved in dH_2_O at a concentration of 1 mg mL^-1^. From this stock solution 50 μL was added to 1 ml of hydrogel precursor solution (0.005 wt%). The hydrogel was polymerized under each Teflon mold and immersed in PBS for 30 minutes to swell out all unreacted Rhodamine dye. Subsequently, the pattern was imaged via confocal microscopy (LSM 780, Zeiss). 3D images were reconstructed from Z-stacks and distants were analyzed using ZEN software (Zeiss).

### 2D and 3D cell culture

#### Cell culture

Human bone marrow-derived stromal cells (hMSCs) were isolated from bone marrow aspirates of healthy donors obtained during orthopaedic procedures with informed consent and in accordance with the local ethical committee (University Hospital Basel; Prof. Kummer; approval date 26/03/2007, Ref. Number 78/07).^78^ Cells were cultured at 37 °C in a humidified atmosphere at 5% CO_2_ in minimal essential medium with alpha modification and nucleosides (MEMα; Sigma-Aldrich) supplemented with fetal bovine serum (FBS, 10%; Gibco), penicillin/streptomycin (P/S, 100 U mL^-1^; Gibco), and fibroblast growth factor 2 (FGF-2, 5 ng mL^-1^; PeproTech). Cells were passaged before reaching 90% confluency and the medium was changed every 2–3 days.

#### Fabrication of hydrogels for 2D cell culture and cell seeding

For 2D cell culture experiments, PEG hydrogels (3.5, 5, and 6 wt%) were prepared with *E* ~ 5, 16, and 35 kPa, respectively. For 3.5 wt% hydrogels, 8-arm PEG-NB (35 mg; 28 mM NB) was mixed with freshly prepared DTT (2.0 mg; 28 mM SH), lithium phenyl-2,4,6-trimethylbenzoylphosphinate (LAP) photoinitiator (2.5 mg; 0.25 mM), CRGDS (1 mM), and dH_2_O. For 5 wt% hydrogels, 8-arm PEG-NB (50 mg; 40 mM NB) was mixed with freshly prepared DTT (2.84 mg; 40 mM SH), LAP (2.5 mg, 0.25 mM), CRGDS (1 mM), and dH_2_O. For 6 wt% hydrogels, 8-arm PEG-NB (50 mg; 48 mM NB) was mixed with freshly prepared DTT (3.4 mg, 48 mM), LAP photoinitiator (2.5 mg, 0.25 mM), CRGDS (1 mM), and dH_2_O.

Two glass slides were separated by a silicon spacer (0.5 mm). One of the glass slides was treated with TMPMA to facilitate adhesion of the hydrogel to the glass surface. The other glass slide was treated with Sigmacote to provide a hydrophobic surface that prevented hydrogel adhesion and enabled hydrogel detachment without damaging the surface. The hydrogel precursor solution was injected between the two glass slides and subsequently polymerized under exposure to UV light (λ = 365 nm; I = 20 mW cm^-2^) for 1 to 3 min depending on the gelation time of the hydrogel determined during rheological testing. After careful removal of the Sigmacote treated glass slide, 4- or 8-well chamber separators (SPL Cell culture slides, SPL Life Sciences Co.) were placed on the hydrogel adhered to the MPTMS treated glass slide. The gel was washed with PBS 3x for 5 min to diffuse out residual LAP, stored in PBS, and sterilized under UV light (l = 254 nm) for 30 min. After that, the gels were immersed in culture medium and stored in the incubator to equilibrate at 37 °C. The cell suspension was then applied in each chamber (20’000 to 30’000 cells cm^-2^) on top of hydrogels and the samples were placed on an orbital shaker for 10 min at 60 rpm. The samples were then stored in the incubator for the further cell culture.

#### Fabrication of hydrogels for 3D cell culture and cell seeding

The hydrogel precursor solution was prepared as described in the procedure for the preparation of 2D hydrogels and additionally supplemented with SAT peptide motif (Ac-GCRE-DDD-LPMTGG-NH_2_; MW = 1406.5 g mol^-1^; 1125 μg mL^-1^; 800 μM). The silicon spacer (0.5 mm) was placed on the TMPMA-treated glass slide and the Teflon mold was used to pattern the hydrogel, as described above. The patterned hydrogel adhered to the TMPMA-treated glass slide was rinsed with PBS 3x for 5 min to diffuse out residual LAP, immersed in PBS, and sterilized under UV light (l = 254 nm) for 30 min.

4- or 8-well chambers (SPL Cell culture slides, SPL Life Sciences Co.) were mounted on the glass slide with the patterned hydrogels. The hydrogel slab was cropped with a sharp spatula to fit the dimensions of the wells in order to secure a proper chamber fixture and avoid leakage. The hydrogel was immersed in culture medium and stored in the incubator to equilibrate at 37 °C. The cells were seeded in 3D niches by gravitational sedimentation. The cell suspension was then applied in each chamber (5’000 to 30’000 cells cm^-2^) on top of patterned hydrogels and the samples were placed on an orbital shaker for 10 min at 60 rpm. The well occupation was examined using a light microscope and, when most niches were occupied by cells, the excess medium was lightly aspirated and samples were gently washed with fresh medium under 60° tilt to remove the cells from the array surface outside of the niches. The samples were stored in the incubator at 37 °C until the niche sealing.

#### Niche sealing and subsequent cell culture

A thin hydrogel sheet (0.1 mm) was prepared, which was used to seal the cell niches after cell sedimentation. The hydrogel was prepared with the same formulation as its patterned equivalent with the exception of a different Sortase A substrate. Sealing hydrogels were prepared with SAG peptide (GGGG-LERCL; Mw = 860.92 g mol^-1^, 551 μg mL^-1^, 800 μM). A 0.1 mm spacer was sandwiched between two glass slides treated with Sigmacote. The hydrogel precursor solution was injected between the glass slides and polymerized upon exposure to UV light (l = 254 nm, I = 20 mW cm^-2^) for 1 to 3 min depending on the hydrogel formulation. The sealing hydrogel layer was rinsed with PBS 3x for min, immersed in PBS, and sterilized under UV light (l = 254 nm) for 30 min. After sterilization, the hydrogel layer was transferred to culture medium and stored in the incubator until further use.

For sealing the niches, the well chambers were removed and Sortase A solution (MW = 21.7 kDa, 20 μL, 86 mg mL^-1^; 4 μM) in Ca^2+^-containing culture medium at 37 °C was applied to the surface of the patterned hydrogel. The thin hydrogel layer containing SAG peptide was placed on top of the patterned substrate and pressed gently, after which the well chambers were again mounted on the glass slide. A minimal amount of culture medium (200 μL) was added prior to storage of the hydrogel in the incubator in order to ensure the covalent crosslinking of the hydrogel slabs without excessive drying. After 1–2 h, 500 μL fresh medium was added to each chamber and the cell-laden arrays were stored in the incubator for culture. The medium was changed every 2 days by substituting half of the old medium with fresh medium.

### Cell function assessment

#### Live/Dead assay

Membrane integrity was assessed after 1 and 3 days of culture using the Live/Dead assay (ThermoFisher). We used membrane integrity as a measure of cell viability in these studies. The gels were incubated in the 4 μM Calcein AM and 2 μM Ethidium homodimer-1 (EthD-1) in PBS. After 30 min of incubation, cell-laden hydrogels were washed twice with PBS for 5 min, resuspended in fresh medium, and imaged.

#### EdU assay

The fraction of proliferating cells was determined by incorporating 5-ethynyl-2’-deoxyuridine (EdU) in the culture medium for a 12 h pulse on the first and third day of culture for both 2D and 3D samples. Incorporation of EdU was visualized by Alexa Fluor 647 azide staining using EdU DetectPro Imaging Kit (BCK-EdUPro-IM647, BaseClick) according to the manufacturer’s protocol.

In brief, hMSCs were seeded on 2D hydrogels and treated with EdU (10 μM) in culture medium 12 h prior to the end of the experiment. After 12 h of EdU treatment, the typical cell cycle period for hMSCs, samples were fixed by treatment with 2% PFA for 15 min and a subsequent treatment with 4% PFA for an additional 15 min. The 2D samples were washed with PBS 3x for 10 min and with PBS supplemented with 5% bovine serum albumin (BSA, Sigma-Aldrich) 2x for 5 min. All samples were permeabilized with TritonX-100 (0.1% in PBS) for 20 min at RT and washed 2x for 5 min with PBS supplemented with 5% BSA.

The 3D samples were treated the same way as 2D samples. The incubation time with DetectPro reaction cocktail containing an azide functionalized Alexa Fluor 657 was extended to 3 h in the dark at RT. The samples were then washed with PBS 3x for 30 min and with PBS supplemented with 5% BSA 2x for 1 h.

In both 2D and 3D samples, the cell nuclei were counterstained with 4’,6-diamidino-2-phenylindole (DAPI; 300 nM; Sigma-Aldrich,) in PBS.

#### YAP staining

2D and 3D hydrogel samples with hMSCs were fixed by treatment with 2% PFA for 15 min and a subsequent treatment with 4% PFA for an additional 15 min. Next, the cells were washed and permeabilized with Triton X-100 (0.5% in PBS). The samples were blocked with 5% BSA in PBS for 1 h and incubated with primary anti-YAP1 antibody (1:300, mouse; Santa Cruz) in 5% BSA and left overnight at 4 °C. The cells were washed 3x for 10 min with PBS with Tween-20 (PBST, 0.05%; Sigma-Aldrich) and the corresponding anti-mouse Alexa Fluor-647 conjugated secondary antibody (1:400, goat; ab150115, Abcam) was applied for 2 h at RT. Nuclei were labelled with DAPI and F-actin was labelled with Phalloidin-iFluor 488 (1:1000, ab176753; Abcam). After the staining, the sample was washed 3x for 10 min with PBST. The sample was resuspended in PBST with P/S and stored in the fridge until imaging.

### Imaging and image analysis

#### Imaging

For assessment of cell function, fluorescence microscopy was performed using an inverted confocal laser scanning microscope (LSM 780, Axio Observer; Zeiss) equipped with an Airyscan detector. The samples were imaged using an EC Plan-Neofluar 10x/ 0.30 Ph1 M27 objective. Each confocal image (1024 × 1024 pixel resolution) was obtained in z-stack (stack depth and step size were chosen according to the niche dimensions). The pixel size was 1.384 x 1.384 μm. All live cell measurements were performed at 37 °C in an atmosphere containing 5% CO_2_. For all 3D samples, the transmitted light channel was recorded for well identification.

For viability investigation, live hMSCs stained with Calcein AM, EthD-1 were illuminated at excitation wavelengths of 494 nm, 528 nm, and 650 nm and detected at 500–525 nm, 600–640 nm, and 655–700 nm, respectively.

For proliferation assessment, fixed hMSCs, stained with DAPI and Alexa Fluor-647 conjugated secondary antibody, were illuminated at excitation wavelengths of 405 nm and 650 nm and detected at 450–480 nm and 655– 700 nm, respectively.

For assessment of YAP localization, fixed hMSCs, stained with DAPI, Phalloidin-iFluor 488, and Alexa Fluor-647 conjugated secondary antibody, were illuminated at excitation wavelengths of 405 nm, 488 nm, and 650 nm and detected at 450–480 nm, 500–550 nm, and 655–700 nm, respectively.

Live imaging was performed on an inverted wide field fluorescence microscope (THUNDER Live Cell; Leica). The samples were imaged using an HC PL Fluorotar 10x/0.32 PH1 objective. Transmitted light channel was recorded over the course of 12 hours at specified positions, the movie was reconstructed from recorded images (**Supplementary Movie S1**). All live cell measurements were performed at 37 °C in an atmosphere containing 5% CO_2_.

For assessment of 3D spreading morphology fluorescence microscopy was performed using a THUNDER Live Cell microscope. The samples were imaged using an HC PL APO 40x/0.95 dry objective. Images were obtained in z-stack (stack depth was chosen according to the niche dimensions and step size was set automatically to optimize ICC). The pictures were processed using THUNDER Large Volume Computational Clearing settings, Feature Scale (nm): 5,000, Strength (%): 92, Adaptive Deconvolution set with a refractive index of the aqueous mounting medium of 1.33. The 3D images were reconstructed from z-stacks and rendered using LAS X 3D software (Leica).

#### Image processing

The images were exported in batch in .lsm or .czi format using ZEN 3.1 blue edition using the z-stack alignment module. The z-stack .lsm or .czi images were exported as TIF RGB images (Live/Dead: maximum intensity projection (MIP), EdU: MIP, YAP: single plane at highest DAPI signal) in batch process using an ImageJ macro. All 3D cell images were overlaid with the AutoCAD masks with the corresponding well pattern to enhance the niche outlines to facilitatie the following analyses. The overlay was performed in MATLAB using 2D crosscorrelation between the binary mask of the well pattern and the binarized bright field image of the wells.

#### Viability analysis

Images were analyzed using Cell Profiler. Separate pipelines were constructed for 2D and 3D experiments. For 2D, the Calcein AM positive cells and EthD-1 positive nuclei were identified using separate IdentifyPrimaryObject modules applying appropriate size and intensity thresholding. The identified objects were related via the RelateObjects module and the fraction of viable cells was reported as the fraction of cells that stained negative for EthD-1.

For 3D, additionally the wells were identified using the IdentifyPrimaryObject module. The objects were related via the RelateObjects module and the wells containing multiple cells or nuclei were excluded from the analysis (only the wells containing single cells were analyzed). The fraction of viable cells was reported as the fraction of encapsulated cells staining negative for EthD-1.

#### Proliferation analysis

Images were analyzed using Cell Profiler. For 2D, the nuclear regions were identified using the IdentifyPrimaryObject module. The fraction of proliferating cells was reported as the fraction of total nuclei that stained positive for EdU divided by the total number of nuclei, indicated by DAPI staining.

For 3D, the wells were identified using the IdentifyPrimaryObject module. The wells with multiple cells were excluded from the analysis (only the wells containing single cells were analyzed).

#### YAP N/C ratio

Images with DAPI and YAP signal were analyzed using Cell Profiler. For 2D, nuclei (DAPI) and YAP^+^ cells were identified using separate IdentifyPrimaryObject modules with appropriate size and intensity thresholds. The nuclear region was identified as a region that stained positive for DAPI. The area outside of the nuclear region that stained positively for YAP was identified as a cytoplasm using the IdentifyTertiaryObject module. YAP mean fluorescence intensities within nuclear and the cytoplasmic regions were calculated using the MeasureObjectIntensity module. The ratio of mean fluorescence intensities within the nucleus and cytoplasm was quantified and reported as the YAP N/C ratio.

For 3D, the wells were identified using the IdentifyPrimaryObject module. The objects were related to each other and only the wells containing single cells were analyzed.

All Cell Profiler pipelines have been uploaded on our GitHub repository: https://github.com/MEL-ETH/Single_Cell_ECM.

### Statistical analysis

All experiments were performed with at least three biological replicates. Statistical analyses were performed with OriginPro 2019. Quantitative data for viability, proliferation, and YAP N/C ratio are presented as mean ± standard error or the mean (s.e.m.). Differences between experimental conditions were assessed using one-way or two-way analysis of variance (ANOVA) with Tukey post-hoc test. The significance levels were set at p = 0.05 (*).

## Acknowledgements

We would like to thank Prof. Marcy Zenobi-Wong and David Fercher for providing us with Sortase A and assisting in synthesis of substrate peptides; Börte Emiroglu for assistance in designing micropatterning methods; ScopeM for assisting us in developing imaging strategies and image analysis pipelines to analyze our data; FIRST-Lab for providing us with the access to clean room facilities; Prof. Christofer Hierold and Prof. Aldo Ferrari for giving us the access to their laboratories and instruments; and Prof. Dr. Ivan Martin at the University Hospital Basel for providing us with hMSCs.

## Author contributions

The project was conceived of and designed by O.D. and M.W.T. The experiments were carried out by O.D., A.B., S.J., M.A.B., N.B., and M.L. A.B., G.B. and N.B. provided assistance in developing image analysis tools or developing the strategy for enzymatic ligation and contributed to writing the supporting information. The manuscript was written by O.D. and M.W.T. and all authors have approved the final version of the manuscript.

## Abbreviations

2D: two-dimensional
3D: three-dimensional
DAPI: 4′, 6-diamidino-2-phenylindole
*E*: Young′s modulus
EdU: 5-ethynyl-2′-deoxyuridine
ECM: extracellular matrix
EthD-1: ethidium homodimer
DTT: dithiothreitol
G′: shear storage modulus
G′′: shear loss modulus
hMSCs: human mesenchymal stem cells
PDMS: polydimethylsiloxane
PEG: poly(ethylene glycol)
YAP: YES-associated protein.

