## Supporting Information for "3D confinement regulates stem cell fate"

### Supplementary Figures

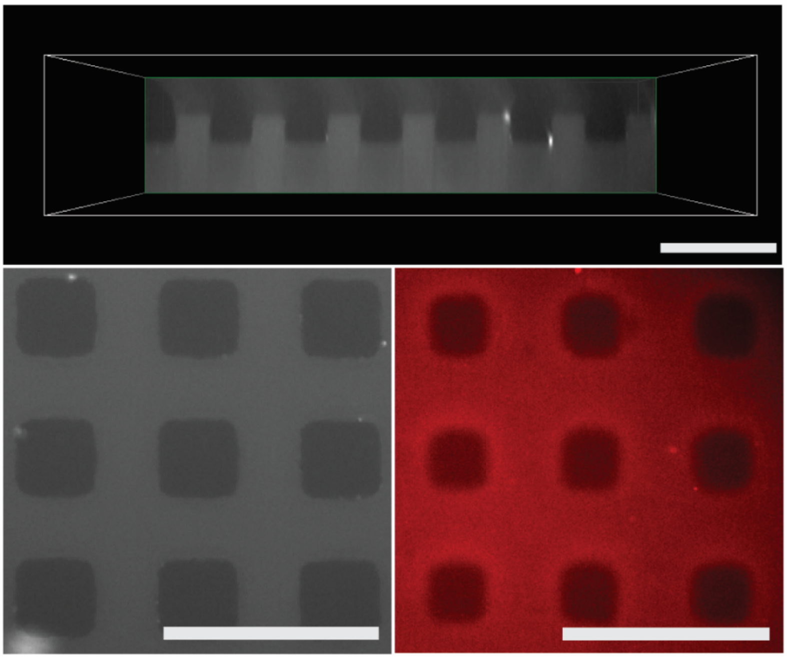

**Figure S1. Teflon molds improved pattern transfer to hydrogels as compared with PDMS molds.** The patterned hydrogel profile was assessed using fluorescence microscopy. Here the hydrogel pattern was imaged in the x-z (top) and x-y (bottom) planes. The hydrogel was fluorescently labeled with acrylated Rhodamine B. The hydrogel casted using Teflon mold (grey) exhibited improved pattern fidelity as compared with hydrogels formed using PDMS molds (red). Scale bars, 150 µm.

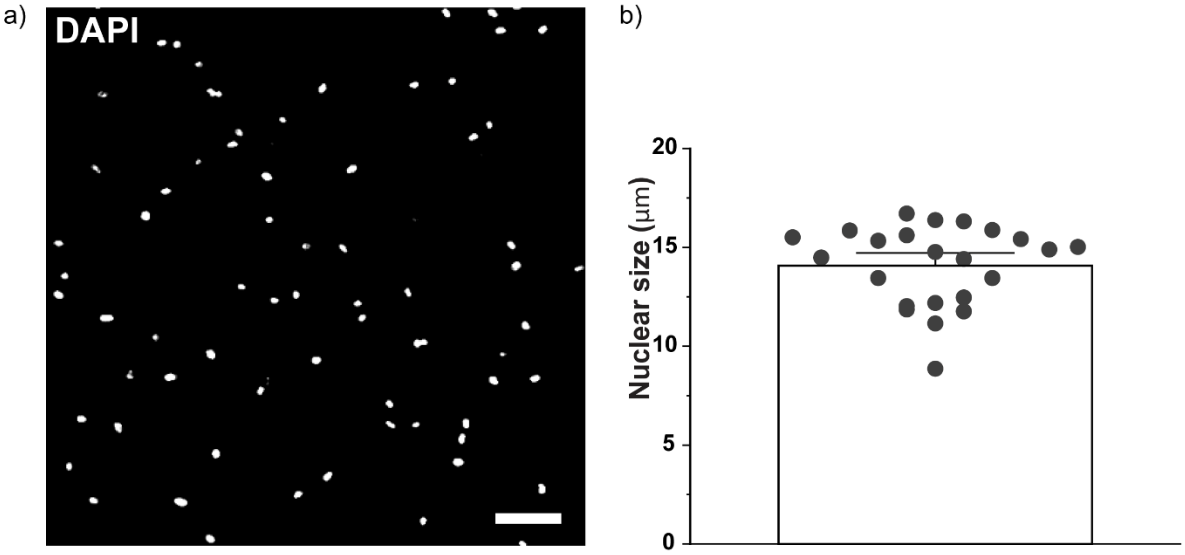

**Figure S2. Size of hMSCs nuclei on stiff 2D hydrogels.** a) The cell cultured on 2D gels were stained with DAPI. b) Nuclear size is represented as the average major length of the nuclear region of hMSCs at medium stiffness 2D substrates (~16 kPa). The length of the major dimension is measured using ImageJ.

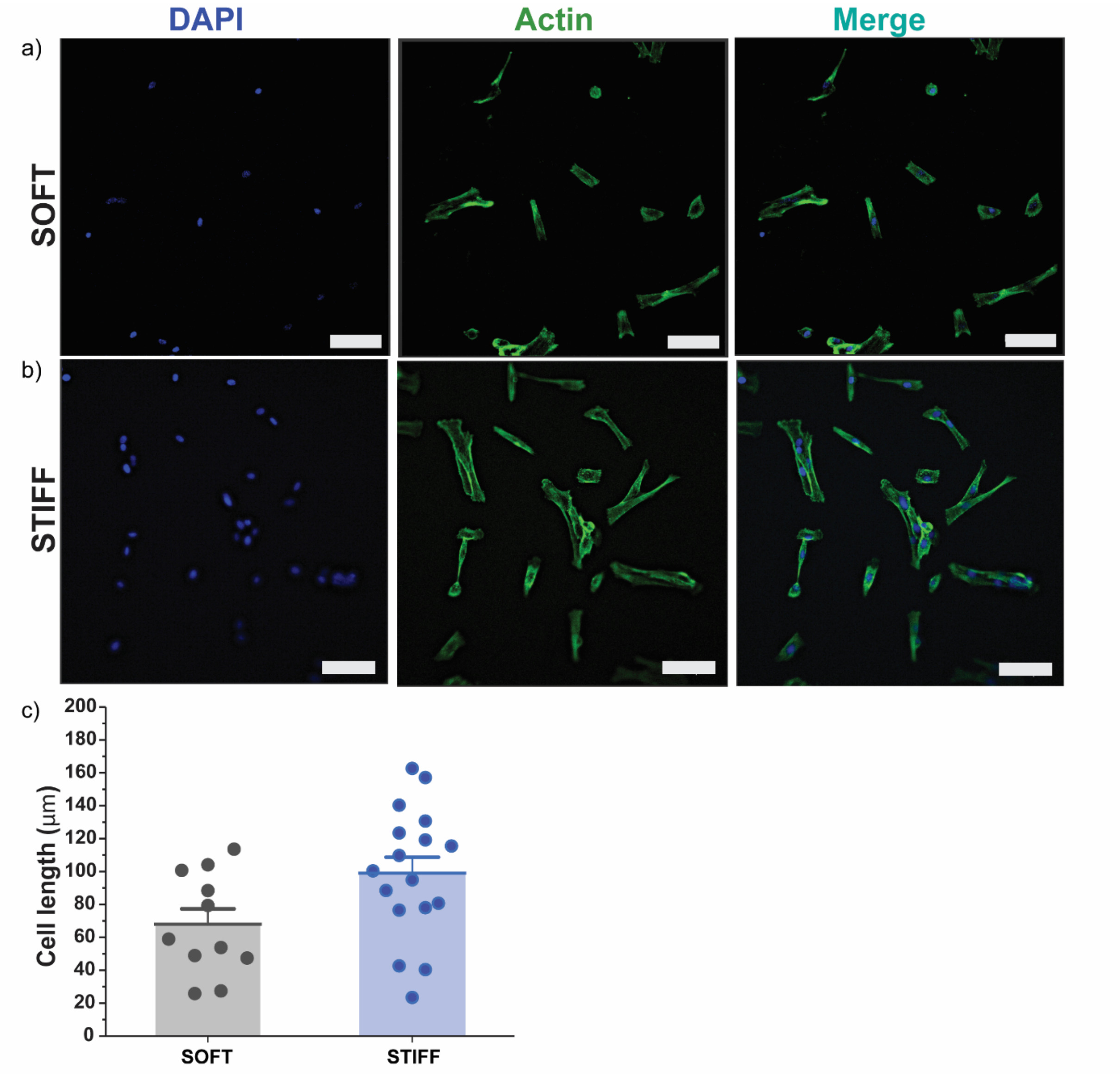

**Figure S3. hMSCs obtained different spread morphologies on soft and stiff hydrogels in 2D.** hMSCs cultured on soft hydrogels appear to be less elongated than on stiff gels, 1 day after seeding. The length of MSCs was calculated using ImageJ. Scale bar, 100 µm.

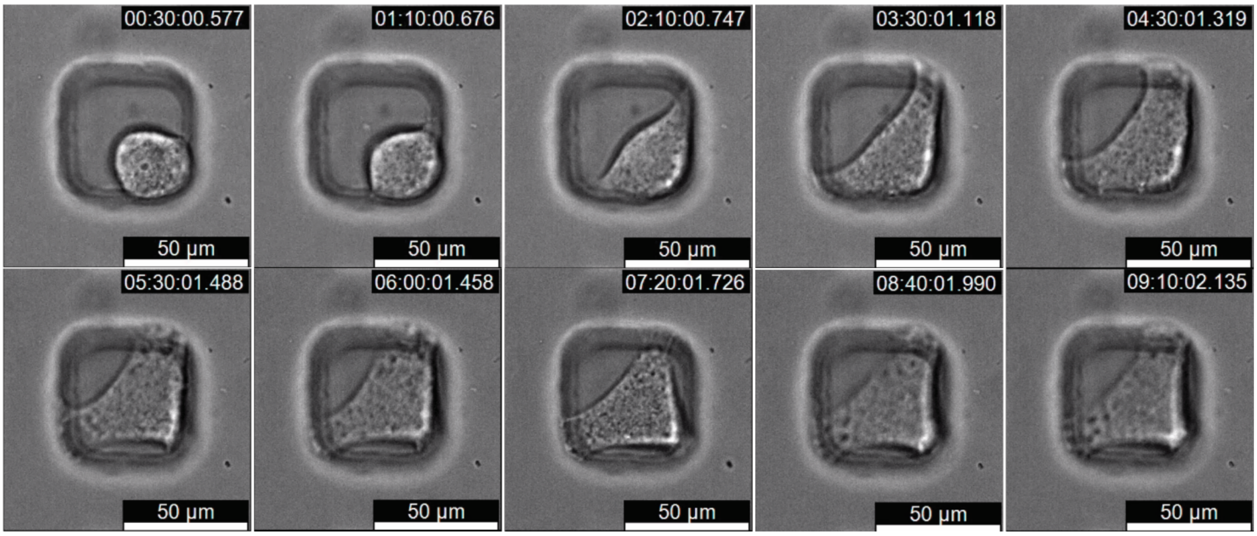

**Figure S4. Cells spread within the 3D microniches within hours after encapsulation.** Live cell imaging (THUNDER Live Cell Microscope, Leica) of hMSCs in V3 niches revealed that the cells spread rapidly after encapsulation. Cell elongation was observed within 2 h after encapsulation. The cell obtained a spread shape following 4–5 h, which remained stable for the remainder of the time course of the imaging experiment. Total elapsed time, ~9 h.

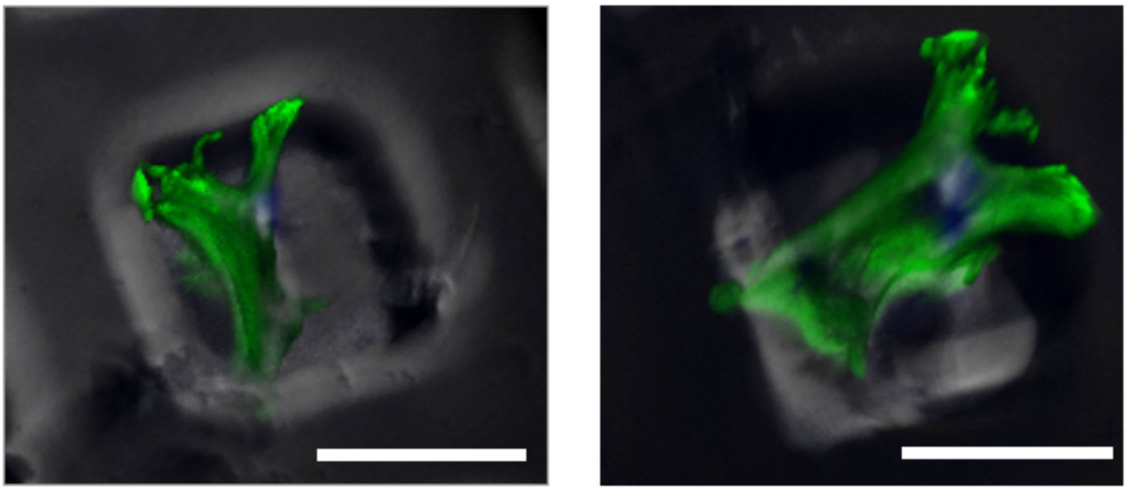

**Figure S5. hMSCs adopted 3D spread morphology in the microniches.** After 3 days, hMSCs spread in 3D within the microniches, as opposed to spreading along the bottom surface of the niche. The image is generated from a 2D stack on Leica THUNDER microscope. Representative images of cells in V3 niches. Actin was labelled with Phalloidin-iFluor 488 (green) and DNA was labelled with DAPI (blue). Scale bar, 50 µm.

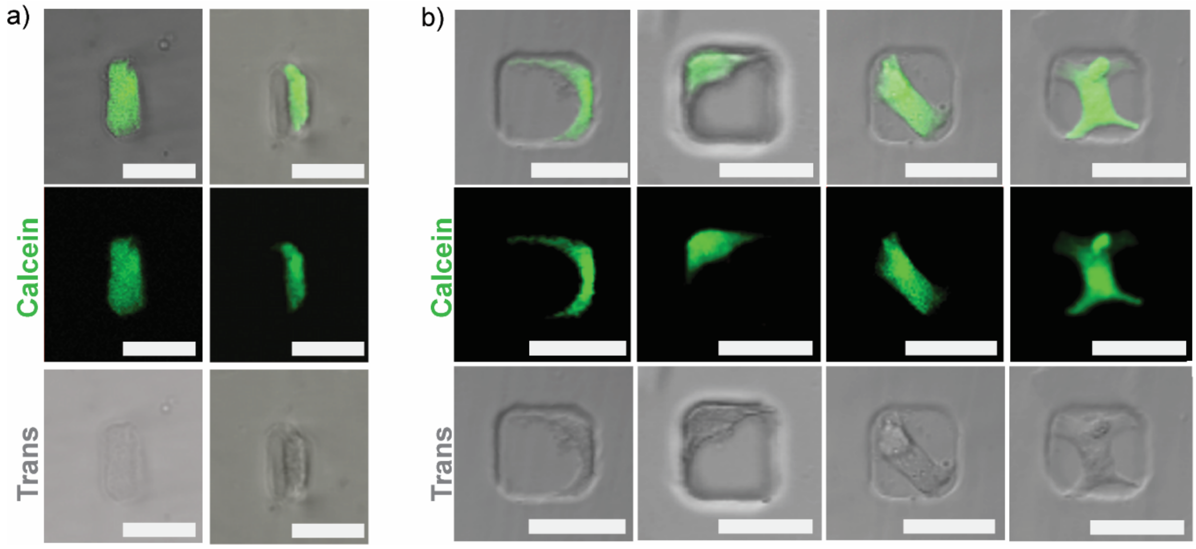

**Figure S6. hMSCs spreading within small V1 and large V3 niches.** a) In V1 niches, the cells are able to spread and occupy most of the niche area. b) In V3 niches, the cells often inhabit only a fraction of the niche space spreading along to the niche walls, in the corners, or diagonally. The cells were stained with Calcein AM (green). Scale bar, 50 µm.

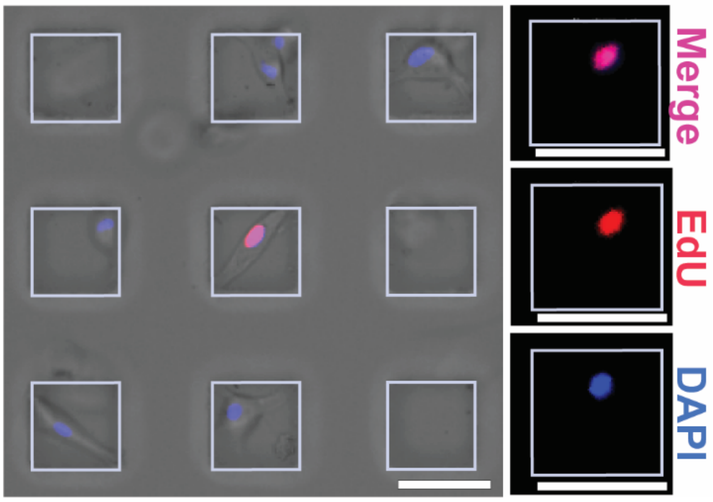

**Figure S7. hMSC proliferation was quantified using EdU staining.** The proliferation was assessed by calculating the fraction of S-phase cells that stained for 5-ethynyl-2′-deoxyuridine (EdU, red). The nuclei were counterstained with 4′,6-diamidino-2-phenylindole (DAPI, blue). Representative image showing single cells cultured within large, V3 microniches. Scale bars, 50 µm.

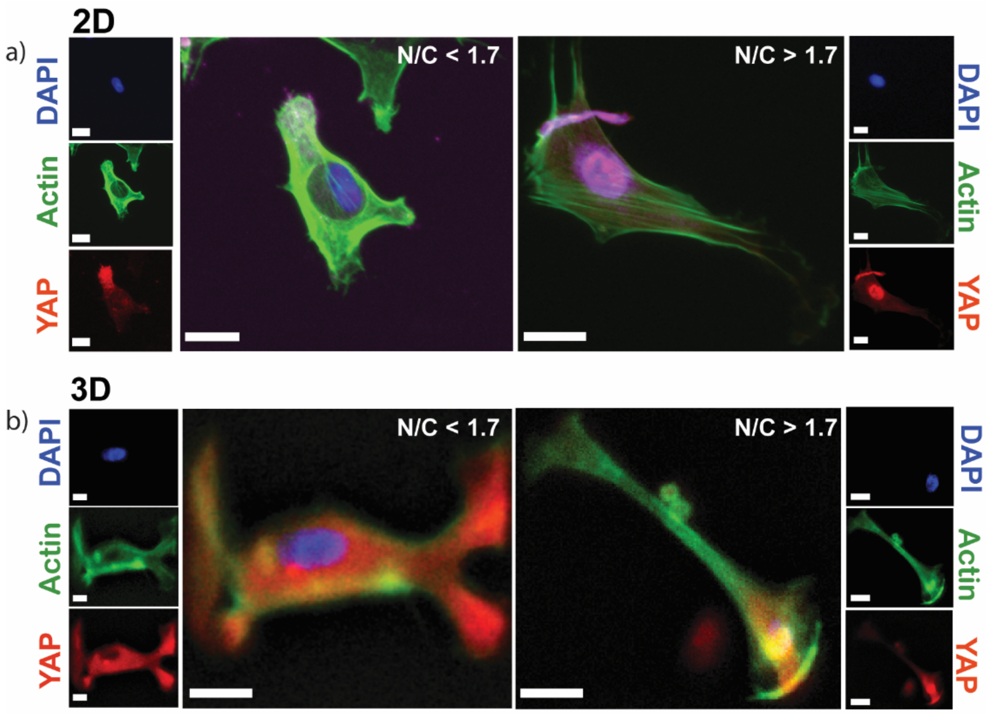

**Figure S8. YAP localization in hMSCs.** YAP localization was assessed using immunocytochemistry in cells cultured on 2D or in 3D hydrogels and defined as NC ratio. The YAP activation is indicated by NC ratio above 1.6. a) hMSCs cultured on 2D gels expressed both cytoplasmic (left) and nuclear (right) YAP localization. Scale bar, 20 (left) and 30 μm (right). b) hMSCS cultured in large V3 microniches also expressed both cytoplasmic (left) and nuclear YAP localization (right). Scale bar, 10 μm.

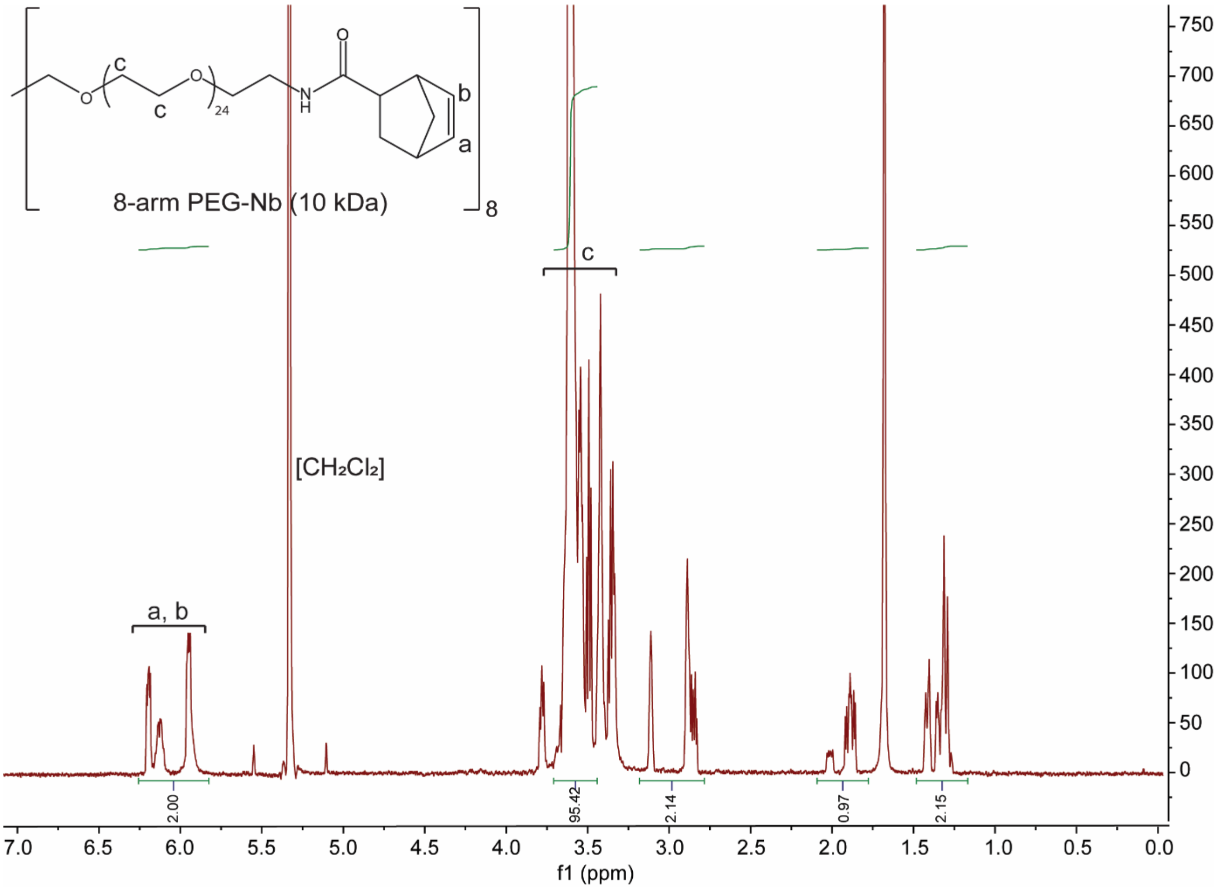

**Figure S9. [1]H NMR spectrum of PEG$_{10kDa}$-norbornene (500MHz, $CD_2Cl_2$).** Functionalization of the 8-arm PEG with norbornene was determined to be above 95% by comparing the integrated area under the peak for the norbornene vinyl protons (δ = 6.0–6.3, m, 2H) and PEG ether protons (δ = 3.5–3.9, m, 96H).

a)

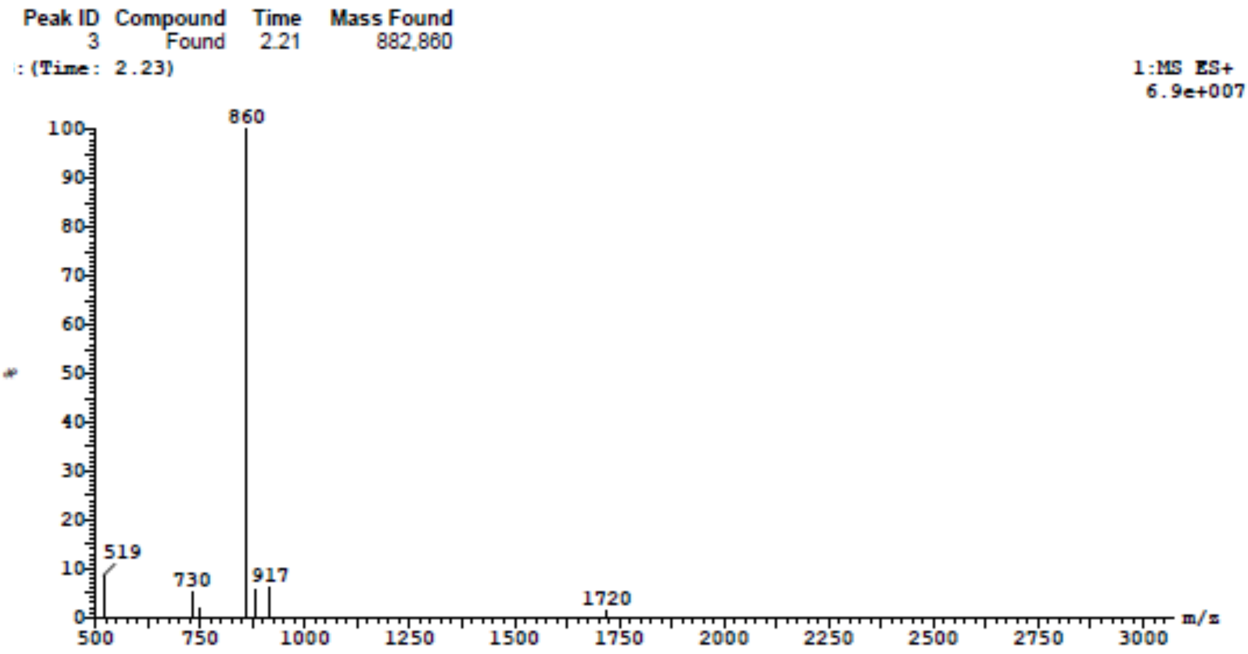

b)

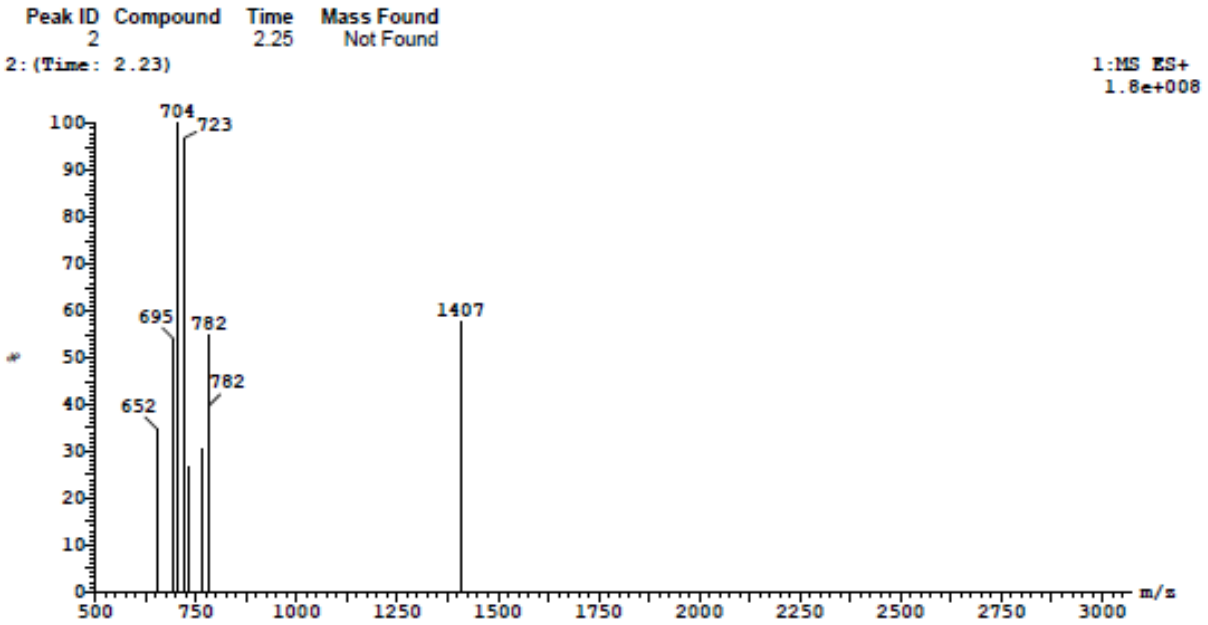

**Figure S10. LC-MS (high resolution, positive mode) spectra of Sortase A substrate peptides SAG (GGGGLERCL) and SAT (Ac-GCRE-DDD-LPMTGG).** a) SAG: calculated mass: 860.98 g $mol^{-1}$; measured m/z: 860 $[M]^+$, 1720 $[2M+H]^+$. b) SAT: calculated mass: 1406.49 g $mol^{-1}$; measured m/z: 704 $[M+2H]^{2+}$, 723 $[M+H+K]^{2+}$, 782 $[M+2K]^{2+}$, 1407 $[M+H]^+$.

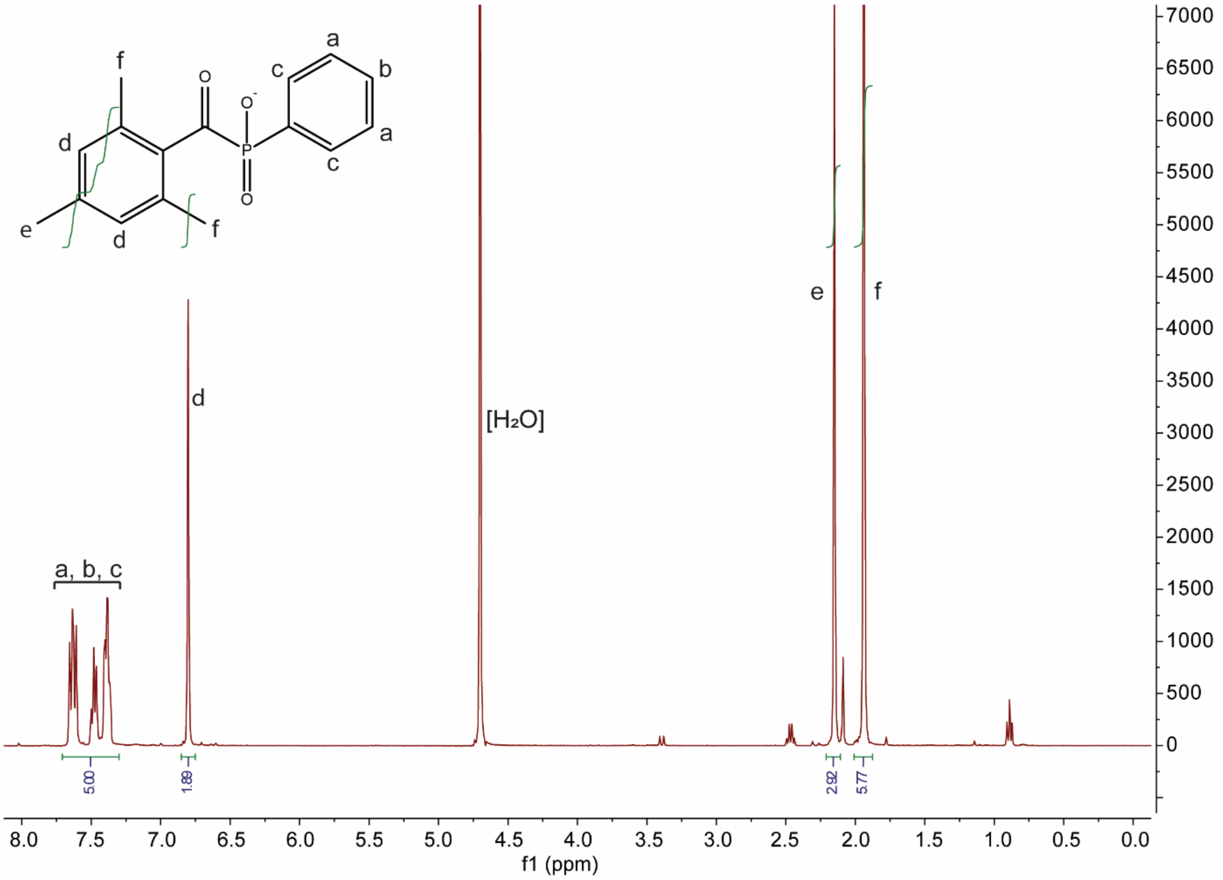

**Figure S11. [1]H NMR spectrum of LAP (500MHz, $D_2O$).** δ 7.75 (m, 2H), 7.59 (m, 1H), 7.49 (m, 2H), 6.91 (s, 2H), 2.26 (s, 3H), 2.05 (s, 6H).

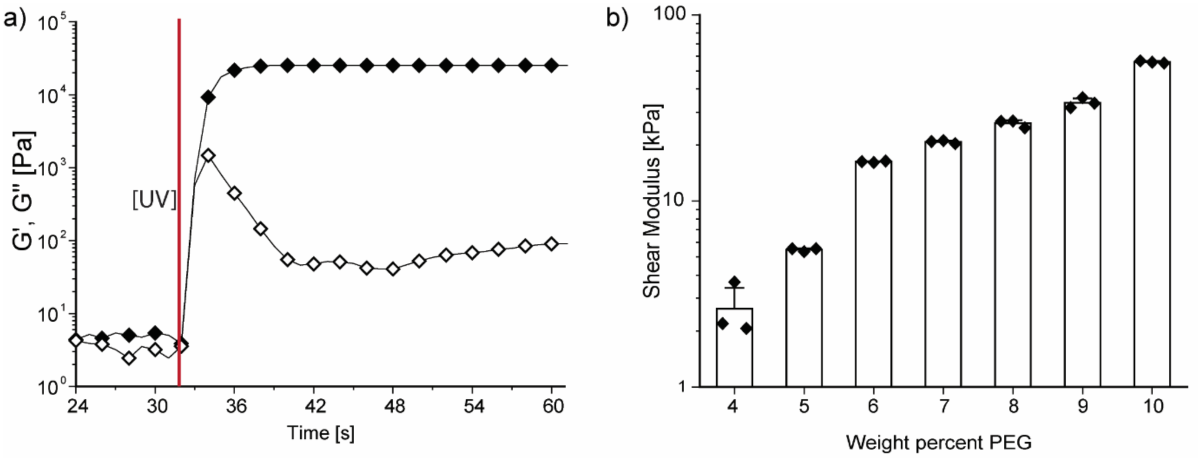

**Figure S12. Mechanical properties of PEG hydrogels.** a) Dynamic time sweep rheology of 8-arm PEG-NB reacted with DTT upon UV light exposure ($\lambda$ = 365 nm, I = 20 mW $cm^{-2}$) demonstrated that gelation occurred within seconds after UV exposure, indicated by the vertical red line. Gelation was complete within 30 s after UV exposure, indicated by G′ reaching a plateau value. b) The equilibrium swollen shear modulus scaled with the PEG wt%, provided a handle to tune the stiffness of the PEG-based hydrogels. Bar plots represent mean + s.e.m.

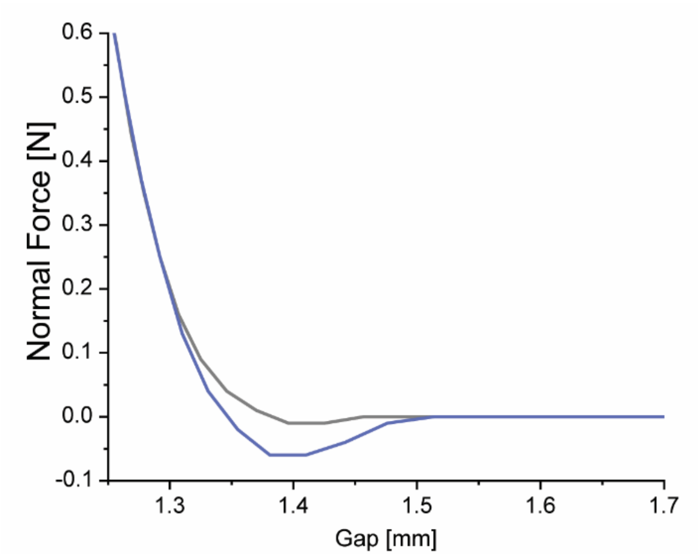

**Figure S13. Normal force ($F_n$)–displacement curves for measuring adhesion between two gels adhered via covalent bond formation between two substrate peptides catalyzed by bacterial enzyme Sortase A.** The pull-off test were performed on the rheometer in extensional mode. The geometry was raised with a velocity of 0.01 mm $s^{-1}$ and $F_n$ upon retraction was measured and $F_n$–displacement curves were recorded. In order to calculate the work of adhesion ($J\,m^{-2}$) from the recorded curves, the retraction force was integrated as a function of the displacement, followed by dividing the resulting adhesion energy by the known contact area (surface area of geometry) at the interface.

### Supplementary Tables

Statistical significance was determined for p-values less than 0.05.

***Cell viability***

*Cell viability 3D Day 1 (Figure 3a)*

After 1 day of culture in 3D niches, hMSCs exhibited high viability without variation in mean viability values between niche volumes or stiffnesses.

Table S1. Two-way ANOVA

| | DF | Sum of Squares | Mean Square | F Value | P Value |
|---|---|---|---|---|---|
| Stiffness | 2 | 69.15756 | 34.57878 | 0.559 | 5.81E-01 |
| Volume | 2 | 42.62572 | 21.31286 | 0.34455 | 0.71311 |
| Interaction | 4 | 482.7966 | 120.6992 | 1.95123 | 0.14552 |
| Model | 8 | 594.5799 | 74.32249 | 1.2015 | 0.35178 |
| Error | 18 | 1113.443 | 61.85794 | -- | -- |

Table S2. Tukey's comparison of means (stiffness)

| | MeanDiff | SEM | q Value | Prob | Alpha | Sig | LCL | UCL |
|---|---|---|---|---|---|---|---|---|
| Medium–Soft | 3.54487 | 3.70759 | 1.35215 | 0.61297 | 0.05 | 0 | -5.91744 | 13.00718 |
| Stiff–Soft | 3.22215 | 3.70759 | 1.22905 | 0.66605 | 0.05 | 0 | -6.24016 | 12.68446 |
| Stiff–Medium | -0.32272 | 3.70759 | 0.1231 | 0.99583 | 0.05 | 0 | -9.78503 | 9.13959 |

Table S3. Tukey's comparison of means (volume)

| | MeanDiff | SEM | q Value | Prob | Alpha | Sig | LCL | UCL |
|---|---|---|---|---|---|---|---|---|
| V2–V1 | 2.7291 | 3.70759 | 1.04098 | 0.74564 | 0.05 | 0 | -6.73321 | 12.19141 |
| V3–V1 | 2.59674 | 3.70759 | 0.9905 | 0.76631 | 0.05 | 0 | -6.86557 | 12.05905 |
| V3–V2 | -0.13235 | 3.70759 | 0.05048 | 0.9993 | 0.05 | 0 | -9.59466 | 9.32996 |

*Viability 3D Day 3 (Figure 3b)*

After 3 days of culture in 3D niches, the mean hMSC viability in high stiffness niches was higher than in low and medium stiffness niches.

Table S4. Two-way ANOVA

| | DF | Sum of Squares | Mean Square | F Value | P Value |
|---|---|---|---|---|---|
| Stiffness | 2 | 2389.653 | 1194.827 | 22.48079 | 7.65E-06 |
| Volume | 2 | 287.7307 | 143.8654 | 2.70684 | 0.09112 |
| Interaction | 4 | 915.791 | 228.9477 | 4.30768 | 0.01126 |
| Model | 8 | 3593.011 | 449.1264 | 8.45036 | 5.48E-05 |
| Error | 20 | 1062.976 | 53.14879 | -- | -- |
| Corrected Total | 28 | 4655.987 | -- | -- | -- |

Table 5. Tukey's comparison of means (stiffness)

| | MeanDiff | SEM | q Value | Prob | Alpha | Sig | LCL | UCL |
|---|---|---|---|---|---|---|---|---|
| Medium–Soft | 7.19691 | 3.34967 | 3.0385 | 0.10529 | 0.05 | 0 | -1.27763 | 15.67145 |
| Stiff–Soft | 21.28969 | 3.26033 | 9.2347 | 6.59E-06 | 0.05 | 1 | 13.04118 | 29.53821 |
| Stiff–Medium | 14.09278 | 3.34967 | 5.9499 | 0.00121 | 0.05 | 1 | 5.61824 | 22.56732 |

Table S6. Tukey's comparison of means (volume)

| | MeanDiff | SEM | q Value | Prob | Alpha | Sig | LCL | UCL |
|---|---|---|---|---|---|---|---|---|
| V2–V1 | 7.35488 | 3.34967 | 3.10519 | 0.09638 | 0.05 | 0 | -1.11966 | 15.82942 |
| V3–V1 | 6.77329 | 3.34967 | 2.85965 | 0.13275 | 0.05 | 0 | -1.70125 | 15.24784 |
| V3–V2 | -0.58159 | 3.26033 | 0.25227 | 0.98263 | 0.05 | 0 | -8.8301 | 7.66693 |

*Cell viability 2D Day 3 (Figure 3c)*

After 3 days of culture on 2D substrates, hMSC viability was close to 100% for all stiffness conditions without significant variation between means.

Table S7. One-way ANOVA

| | DF | Sum of Squares | Mean Square | F Value | Prob>F |
|---|---|---|---|---|---|
| Model | 2 | 2.44186 | 1.22093 | 0.31901 | 0.73848 |
| Error | 6 | 22.96363 | 3.82727 | | |
| Total | 8 | 25.40548 | | | |

Table S8. Tukey's comparison of means (stiffness)

| | MeanDiff | SEM | q Value | Prob | Alpha | Sig | LCL | UCL |
|---|---|---|---|---|---|---|---|---|
| Medium Low | -1.24383 | 1.59735 | 1.10123 | 0.7287 | 0.05 | 0 | -6.14505 | 3.65738 |
| High Low | -0.86806 | 1.59735 | 0.76854 | 0.85348 | 0.05 | 0 | -5.76927 | 4.03315 |
| High Medium | 0.37577 | 1.59735 | 0.33269 | 0.9701 | 0.05 | 0 | -4.52544 | 5.27698 |

***Proliferation***

*Proliferation 3D Day 1 (Figure 4a)*

After 1 day in 3D culture, hMSC proliferation was low and was not significantly affected by niche stiffness or volume.

Table S9. Two-way ANOVA

| | DF | Sum of Squares | Mean Square | F Value | P Value |
|---|---|---|---|---|---|
| Stiffness | 2 | 22.06112 | 11.03056 | 2.4523 | 0.11433 |
| Volume | 2 | 0.35833 | 0.17916 | 0.03983 | 0.96104 |
| Interaction | 4 | 5.38363 | 1.34591 | 0.29922 | 0.87461 |
| Model | 8 | 27.80308 | 3.47539 | 0.77264 | 0.63132 |
| Error | 18 | 80.96499 | 4.49806 | -- | -- |
| Corrected Total | 26 | 108.7681 | -- | -- | -- |

Table S10. Tukey's comparison of means (stiffness)

| | MeanDiff | SEM | q Value | Prob | Alpha | Sig | LCL | UCL |
|---|---|---|---|---|---|---|---|---|
| Medium–Soft | -2.17097 | 0.99978 | 3.07088 | 0.10359 | 0.05 | 0 | -4.72256 | 0.38063 |
| Stiff–Soft | -0.70862 | 0.99978 | 1.00236 | 0.76148 | 0.05 | 0 | -3.26022 | 1.84297 |
| Stiff–Medium | 1.46235 | 0.99978 | 2.06852 | 0.33159 | 0.05 | 0 | -1.08925 | 4.01394 |

Table S11. Tukey's comparison of means (volume)

| | MeanDiff | SEM | q Value | Prob | Alpha | Sig | LCL | UCL |
|---|---|---|---|---|---|---|---|---|
| V2–V1 | 0.23025 | 0.99978 | 0.32569 | 0.97123 | 0.05 | 0 | -2.32135 | 2.78184 |
| V3–V1 | 0.25641 | 0.99978 | 0.36269 | 0.96446 | 0.05 | 0 | -2.29519 | 2.808 |
| V3–V2 | 0.02616 | 0.99978 | 0.03701 | 0.99962 | 0.05 | 0 | -2.52543 | 2.57776 |

*Proliferation 3D Day 3 (Figure 4b)*

After 3 days in 3D niches, the proliferation rate settled below 10% and mean proliferation was significantly affected by both stiffness and volume.

Table S12. Two-way ANOVA

| | DF | Sum of Squares | Mean Square | F Value | P Value |
|---|---|---|---|---|---|
| Stiffness | 2 | 31.09483 | 15.54742 | 6.03209 | 0.00989 |
| Volume | 2 | 34.54075 | 17.27038 | 6.70056 | 0.00668 |
| Interaction | 4 | 11.18792 | 2.79698 | 1.08517 | 0.39328 |
| Model | 8 | 76.8235 | 9.60294 | 3.72575 | 0.00975 |
| Error | 18 | 46.39415 | 2.57745 | -- | -- |
| Corrected Total | 26 | 123.2177 | -- | -- | -- |

Table S13. Tukey's comparison of means (stiffness)

| | MeanDiff | SEM | q Value | Prob | Alpha | Sig | LCL | UCL |
|---|---|---|---|---|---|---|---|---|

| | | | | | | | | |
|---|---|---|---|---|---|---|---|---|
| Medium–Soft | 2.01741 | 0.75681 | 3.76982 | 0.03984 | 0.05 | 1 | 0.08591 | 3.94891 |
| Stiff–Soft | 2.46816 | 0.75681 | 4.61211 | 0.01149 | 0.05 | 1 | 0.53666 | 4.39966 |
| Stiff–Medium | 0.45075 | 0.75681 | 0.84229 | 0.82422 | 0.05 | 0 | -1.48075 | 2.38225 |

Table S14. Tukey's comparison of means (volume)

| | MeanDiff | SEM | q Value | Prob | Alpha | Sig | LCL | UCL |
|---|---|---|---|---|---|---|---|---|
| V2–V1 | 1.85184 | 0.75681 | 3.46042 | 0.06149 | 0.05 | 0 | -0.07966 | 3.78334 |
| V3–V1 | 2.71052 | 0.75681 | 5.065 | 0.00575 | 0.05 | 1 | 0.77902 | 4.64202 |
| V3–V2 | 0.85869 | 0.75681 | 1.60458 | 0.50594 | 0.05 | 0 | -1.07281 | 2.79019 |

*Proliferation 2D Day 1 (Figure 4c)*

The mean proliferation of hMSCs increased with stiffness of the 2D substrates, the variation between mean proliferation values of hMSCs on low and high stiffness substrates was significant.

Table S15. One-way ANOVA

| | DF | Sum of Squares | Mean Square | F Value | Prob>F |
|---|---|---|---|---|---|
| Model | 2 | 411.2374 | 205.6187 | 5.14752 | 0.04992 |
| Error | 6 | 239.671 | 39.94517 | | |
| Total | 8 | 650.9084 | | | |

Table S16. Tukey's comparison of means (stiffness)

| | MeanDiff | SEM | q Value | Prob | Alpha | Sig | LCL | UCL |
|---|---|---|---|---|---|---|---|---|
| Medium–Low | 5.46792 | 5.16044 | 1.49848 | 0.5704 | 0.05 | 0 | -10.3661 | 21.30194 |
| High– Low | 16.26892 | 5.16044 | 4.45848 | 0.04509 | 0.05 | 1 | 0.4349 | 32.10294 |
| High–Medium | 10.801 | 5.16044 | 2.96 | 0.17151 | 0.05 | 0 | -5.03302 | 26.63502 |

*Proliferation 2D Day 3 (Figure 4d)*

After 3 days on 2D gels, mean hMSC proliferation increased with stiffness. The differences in mean proliferation values of hMSCs between low and high and medium and high stiffness substrates were significant.

Table S17. One-way ANOVA

| | DF | Sum of Squares | Mean Square | F Value | Prob>F |
|---|---|---|---|---|---|
| Model | 2 | 621.3567 | 310.6784 | 8.24585 | 0.01444 |
| Error | 7 | 263.7386 | 37.67694 | | |
| Total | 9 | 885.0953 | | | |

Table S18. Tukey's comparison of means (stiffness)

| | MeanDiff | SEM | q Value | Prob | Alpha | Sig | LCL | UCL |
|---|---|---|---|---|---|---|---|---|
| Medium–Low | 1.0108 | 1.23004 | 1.16215 | 0.7044 | 0.05 | 0 | -2.76339 | 4.78499 |

| | | | | | | | | |
|---|---|---|---|---|---|---|---|---|
| High– Low | 14.27127 | 1.23004 | 16.4081 | 6.09E-05 | 0.05 | 1 | 10.49708 | 18.04546 |
| High–Medium | 13.26047 | 1.23004 | 15.24595 | 9.29E-05 | 0.05 | 1 | 9.48627 | 17.03466 |

***YAP localization***

*YAP N/C 2D Day 3 (Figure 5b)*

After 3 days on 2D culture, the N/C ratio increased with substrate stiffness. The mean N/C ratio values between medium and low stiffness substrates were significantly different.

Table S19. One-way ANOVA

| | DF | Sum of Squares | Mean Square | F Value | Prob>F |
|---|---|---|---|---|---|
| Model | 2 | 0.45691 | 0.22845 | 8.88094 | 0.0161 |
| Error | 6 | 0.15434 | 0.02572 | | |
| Total | 8 | 0.61125 | | | |

Table S20. Tukey's comparison of means (stiffness)

| | MeanDiff | SEM | q Value | Prob | Alpha | Sig | LCL | UCL |
|---|---|---|---|---|---|---|---|---|
| Medium–Low | 0.55179 | 0.13096 | 5.95892 | 0.01324 | 0.05 | 1 | 0.14998 | 0.95361 |
| High–Low | 0.28574 | 0.13096 | 3.08581 | 0.15316 | 0.05 | 0 | -0.11607 | 0.68756 |
| High–Medium | -0.26605 | 0.13096 | 2.87311 | 0.1854 | 0.05 | 0 | -0.66786 | 0.13577 |

*YAP percent activation Day 3 (Figure 5c)*

After 3 days in 3D niches, mean YAP activation (defined as N/C > 1.7) did not correlate with niche volume. A weak dependence with stiffness was observed, with highest variation between high and medium stiffnesses.

Table S21. Two-way ANOVA

| | DF | Sum of Squares | Mean Square | F Value | P Value |
|---|---|---|---|---|---|
| Stiffness | 2 | 0.06332 | 0.03166 | 3.5171 | 0.05136 |
| Volume | 2 | 0.03133 | 0.01566 | 1.74 | 0.20377 |
| Interaction | 4 | 0.01868 | 0.00467 | 0.51881 | 0.72301 |
| Model | 8 | 0.11333 | 0.01417 | 1.57368 | 0.20156 |
| Error | 18 | 0.16203 | 0.009 | -- | -- |
| Corrected Total | 26 | 0.27536 | -- | -- | -- |

Table S22. Tukey's comparison of means (stiffness)

| | MeanDiff | SEM | q Value | Prob | Alpha | Sig | LCL | UCL |
|---|---|---|---|---|---|---|---|---|
| Medium–Low | -0.02303 | 0.04473 | 0.72829 | 0.86507 | 0.05 | 0 | -0.13718 | 0.09111 |
| High–Low | 0.08926 | 0.04473 | 2.82231 | 0.14196 | 0.05 | 0 | -0.02489 | 0.2034 |
| High–Medium | 0.11229 | 0.04473 | 3.5506 | 0.05427 | 0.05 | 0 | -0.00186 | 0.22644 |

Table S23. Tukey's comparison of means (volume)

| | MeanDiff | SEM | q Value | Prob | Alpha | Sig | LCL | UCL |
|---|---|---|---|---|---|---|---|---|
| V2–V1 | 0.00866 | 0.04473 | 0.27394 | 0.97955 | 0.05 | 0 | -0.10548 | 0.12281 |
| V3–V1 | 0.0762 | 0.04473 | 2.40935 | 0.23101 | 0.05 | 0 | -0.03795 | 0.19034 |
| V3–V2 | 0.06753 | 0.04473 | 2.13541 | 0.30987 | 0.05 | 0 | -0.04661 | 0.18168 |

*YAP N/C ratio 3D Day 1 (Figure 5d)*

On day 1 in 3D niches, mean YAP N/C ratios did not correlate with niche volume and exhibited a weak trend with stiffness, specifically N/C values between low and high stiffness niches were significantly different.

Table S24. Two-way ANOVA

| | DF | Sum of Squares | Mean Square | F Value | P Value |
|---|---|---|---|---|---|
| Stiffness | 2 | 0.67774 | 0.33887 | 5.35206 | 0.01197 |
| Volume | 2 | 0.02955 | 0.01478 | 0.23338 | 0.79363 |
| Model | 4 | 0.69831 | 0.17458 | 2.75727 | 0.05113 |
| Error | 24 | 1.51957 | 0.06332 | -- | -- |
| Corrected Total | 28 | 2.21788 | -- | -- | -- |

Table S25. Tukey's comparison of means (stiffness)

| | MeanDiff | SEM | q Value | Prob | Alpha | Sig | LCL | UCL |
|---|---|---|---|---|---|---|---|---|
| Medium–Low | 0.12437 | 0.11862 | 1.48286 | 0.55434 | 0.05 | 0 | -0.17184 | 0.42059 |
| High–Low | 0.35842 | 0.1131 | 4.48183 | 0.01112 | 0.05 | 1 | 0.07599 | 0.64085 |
| High–Medium | 0.23405 | 0.1131 | 2.9266 | 0.11765 | 0.05 | 0 | -0.04839 | 0.51648 |

Table S26. Tukey's comparison of means (volume)

| | MeanDiff | SEM | q Value | Prob | Alpha | Sig | LCL | UCL |
|---|---|---|---|---|---|---|---|---|
| V2 –V1 | -0.00194 | 0.11561 | 0.02374 | 0.99984 | 0.05 | 0 | -0.29066 | 0.28678 |
| V3 –V1 | -0.05693 | 0.11253 | 0.71552 | 0.86916 | 0.05 | 0 | -0.33795 | 0.22408 |
| V3 –V2 | -0.05499 | 0.11561 | 0.67269 | 0.88335 | 0.05 | 0 | -0.34371 | 0.23373 |

YAP N/C ratio 3D Day 3 (Figure 5e)

After 3 days in 3D niches, mean YAP N/C ratios exhibited a weak correlation with stiffness and volume but were not significantly different.

Table S27. Two-way ANOVA

| | DF | Sum of Squares | Mean Square | F Value | P Value |
|---|---|---|---|---|---|
| Stiffness | 2 | 0.15858 | 0.07929 | 2.42836 | 0.11145 |
| Volume | 2 | 0.1548 | 0.0774 | 2.37056 | 0.11686 |
| Model | 4 | 0.31338 | 0.07835 | 2.39946 | 0.08092 |
| Error | 22 | 0.71833 | 0.03265 | -- | -- |

| | | | | | |
|---|---|---|---|---|---|
| Corrected Total | 26 | 1.03172 | -- | -- | -- |

Table S28. Tukey's comparison of means (stiffness)

| | MeanDiff | SEM | q Value | Prob | Alpha | Sig | LCL | UCL |
|---|---|---|---|---|---|---|---|---|
| Medium Low | 0.06709 | 0.08518 | 1.11387 | 0.71437 | 0.05 | 0 | -0.14689 | 0.28107 |
| High Low | 0.18538 | 0.08518 | 3.07776 | 0.09774 | 0.05 | 0 | -0.0286 | 0.39936 |
| High Medium | 0.11829 | 0.08518 | 1.96389 | 0.36383 | 0.05 | 0 | -0.09569 | 0.33227 |

Table S29. Tukey's comparison of means (volume)

| | MeanDiff | SEM | q Value | Prob | Alpha | Sig | LCL | UCL |
|---|---|---|---|---|---|---|---|---|
| V2 V1 | 0.04899 | 0.08518 | 0.81329 | 0.83468 | 0.05 | 0 | -0.16499 | 0.26297 |
| V3 V1 | 0.17942 | 0.08518 | 2.97873 | 0.11161 | 0.05 | 0 | -0.03456 | 0.3934 |
| V3 V2 | 0.13043 | 0.08518 | 2.16543 | 0.2962 | 0.05 | 0 | -0.08355 | 0.34441 |

### Supplementary Movies

**Supplementary Movie 1. Sortase A sealing**. Demonstrates the adhesion between two hydrogel slabs following enzymatic ligation between two substrate peptides of bacterial enzyme Sortase A. The adhesion between two gels holds under agitation.

**Supplementary Movie 2. Live cells**. This movie follows a single hMSC spreading in large V3 niche for 9 h post seeding.

**Supplementary Movie 3. 3D Cell Spreading**. A short video of a single hMSC spread in 3D within large V3 niche (**Figure S5**) showing the cell from different perspectives.
